# Regulatory T cells crosstalk with tumor and endothelium through lymphotoxin signaling

**DOI:** 10.1101/2024.10.01.616077

**Authors:** Wenji Piao, Long Wu, Yanbao Xiong, Gregory C. Zapas, Christina M. Paluskievicz, Robert S. Oakes, Sarah M. Pettit, Jessica Schmitz, Philipp Ivanyi, Amol C. Shetty, Yang Song, Dejun Kong, Margaret Sleeth, Keli L. Hippen, Young Lee, Lushen Li, Marina W. Shirkey, Allison Kensiski, Aamna Alvi, Kevin Ho, Vikas Saxena, Jan H. Bräsen, Christopher M. Jewell, Bruce R. Blazar, Reza Abdi, Jonathan S. Bromberg

## Abstract

Regulatory T cells (Tregs) are suppressors of anti-tumor immunity that exert multifaceted functions by signaling surrounding cells. We revealed Tregs use their high-level surface lymphotoxin (LT)α1β2 to preferentially stimulate LTβ receptor (LTβR) nonclassical NFκB signaling on both tumor and lymphatic endothelial cells (LECs) to accelerate tumor growth and metastasis. Selectively targeting LTβR nonclassical NFκB pathways on both tumors and LECs cocultured with Tregs, inhibited tumor growth and migration in vitro. Further, we identified protumorigenic chemokines and interferon-stimulated response genes selectively driven by LTβR nonclassical NFκB in melanoma cells. Endothelial specific genes related to oncogenic process such as SOX18 and FLRT2 were identified to be driven under LTβR nonclassical NFκB in LECs. Leveraging *in vivo* Treg LTα1β2 interactions with LTβR on tumor and LECs, transfer of WT but not LTα-deficient Tregs promoted transplanted WT B16F10 growth and tumor cell-derived CXCL1 and CXCL10 secretion in LTβR-deficient host mice, and increased endothelial specific genes related to tumor angiogenesis and lymphangiogenesis, in WT mice bearing LTβR-depleted melanoma. Selectively blocking LTβR nonclassical NFκB pathways remarkably suppressed tumor growth and lymphatic metastasis by reducing tumor cell and LEC-derived CXCL1 and CXCL10 production, restricting Treg and myeloid-derived suppressor cell (MDSC) recruitment to tumor. It also retained intratumoral effector T cells, especially IFNγ^+^ CD8 T cells by restraining Treg facilitated lymphatic vessel permeability. Our data revealed that Treg LTα1β2 promotes LTβR nonclassical NFκB signaling in tumor cells and LECs providing a rational strategy to modulate Treg-mediated protumorigenic molecules to prevent tumor growth and metastasis.

## INTRODUCTION

Lymphotoxin beta receptor (LTβR) is a member of the tumor necrosis factor receptor (TNFR) family and mediates unique immune functions critical for the development and maintenance of various lymphoid microenvironments, including stromal cell specialization, positioning of lymphocytes within lymph nodes (LNs), specialization of high endothelial venules, and formation of secondary lymphatic organs and tertiary lymphoid structure organs^1^. LTβR has two known ligands: the membrane bound lymphotoxin (LT) heterotrimer (LTα1β2), and LIGHT (TNFSF14)^2^. Unlike TNFRI that exclusively signals through the classical nuclear factor-κB (NFκB) pathway, LTβR signals through a dual cascade that activates both classical and nonclassical NFκB pathways in the lymphatic endothelium^2^. The classical pathway acts rapidly, phosphorylating the inhibitor of kappa-B kinase (IKK) complex and degrading the inhibitor IκBα, thereby allowing for release of the RelA/p50 complex and nuclear translocation. The nonclassical pathway has a more sustained response involving the NFκB inducing kinase (NIK) dependent processing of p100 to p52, which dimerizes with RelB for nuclear translocation ^3^.

While the LT system has a defined role in several important aspects of immune regulation and interactions of leukocytes and blood vascular and lymphatic endothelium, there is conflicting evidence for the role LTβR plays in cancer cell biology. Some reports describe links of LTβR to primary tumorigenesis related to mechanisms controlling cell survival, cell-cycle progression, and initiation of angiogenesis^4^. In multiple myeloma, mutations resulting in constitutive activation of the NIK pathway are frequently observed along with associated overexpression of LTβR^5^. LTβR expressing fibrosarcoma cells have been shown to interact with LTα1β2 expressing lymphocytes, resulting in activation of a proangiogenic pathway required for solid tumor neovascularization^6^. In contrast, other reports demonstrated LTβR activation by LIGHT promoted anti-tumor immunity^7^. Agonist anti-LTβR monoclonal antibody (mAb) treatment inhibited tumor growth in colorectal cancers^8^ and enhanced chemotherapeutic responses^1^. Thus, LTβR signaling has a more complex role in the tumor microenvironment (TME) that warrants further investigation.

High frequencies of immunosuppressive regulatory T cells (Tregs) are present in the TME of various primary tumors, and these intratumoral Tregs are key suppressors of anti-tumor immunity^9,10^. Selectively targeting Tregs in the TME has emerged as an effective anti-tumor strategy. In our previous studies, we found that among T cell subsets, both human and mouse Tregs express the highest level of membrane anchored LTα1β2, suggesting that Tregs are the predominant population to interact with LTβR, which is highly expressed on human and mouse LECs^11,12^. LTα deficiency prevents Treg lymphatic migration to draining LNs and impairs their suppressive function in models of allograft protection^11^. Engagement of LTβR on LECs by LTα1β2^+^ but not LTα^−/−^ Treg cells induces LEC basal protrusions that support Treg afferent transendothelial migration (TEM)^11^. Blood endothelial cells including high endothelial venules (HEV) also express high levels of LTβR. However, the LTαβ-LTβR interaction is not required for Treg cell migration from blood through HEV into the LNs^11^, although deletion of endothelial LTβR in mice impairs LN homing of conventional T cells and B cells through impaired integrin and selectin expression by HEV^13^. Since tumor cells also express LTβR, Tregs may also have a direct impact on tumors through LTαβ/LTβR pathways.

While LTα1β2-LTβR interactions are integral to many aspects of immune cell migration and endothelial cell responses, there are limited studies that address the roles that LTβR may play in cancer cell migration, interactions with blood or lymphatic endothelium, and initiation of metastasis. Previous studies^1,8,14^ assessing the role of LTβR in cancer have used approaches that relied on total receptor blockade or activation with pharmacologic or genetic interventions but did not dissect the contributions of each bifurcated LTβR signaling cascade to specific cell processes, including migration patterns. A more in-depth understanding can be obtained only with selective targeting of the LTβR-mediated classical or nonclassical NFκB signaling pathways. In our previous studies, we constructed and validated novel cell permeable decoy receptor peptides based on the LTβR cytoplasmic TRAF recruitment domains^3^. These peptides were capable of selectively and specifically targeting each arm of the LTβR NFκB signaling pathways.

Since most of solid tumors contain both tumor and stroma cells, including LECs which form lymphatic vessels for transporting metastatic and immune cells to draining LNs, We focused on LTβR signaling in tumor cells and LECs. We used these LTβR decoy peptides to selectively target either classical or nonclassical NFκB signaling pathways in melanoma cells and LECs to dissect the molecular underpinnings related to cancer cell migration and metastasis in vitro and in vivo syngeneic mouse melanoma model whose metastasis follow reliable lymphatic routes^15^. Our results demonstrate that selective inhibition of tumor and LEC LTβR nonclassical NFκB signaling pathway significantly suppresses human and mouse melanoma and other cancer cell growth and TEM in vitro. When applied to an in vivo mouse model of melanoma metastases, the LTβR nonclassical NFκB blocking peptides effectively limit tumor cell and immune cell lymphatic migration into the tumor draining LNs and induce significant tumor regression. Mechanistically, constitutively activated tumor LTβR nonclassical NFκB signaling harnesses chemokines and interferon-regulated genes which promote immune suppressive cell recruitment, especially Tregs which further enhance LTβR nonclassical NFκB signaling in tumor and LECs. These results indicate a general reliance on LTβR signaling in cancer cell migration and the initiation of lymphatic based metastases that could be utilized for novel therapeutic modalities.

## RESULTS

### High LTβR expression on tumors denotes worse patient prognosis

To assess LTβR expression and its association with phenotype and clinical characteristics in human cancer samples, The Cancer Genome Atlas (TCGA) gene expression data and the Human Protein Atlas were queried for LTβR expression levels and patient survival. High LTβR expression on tumors was associated with poor survival in patients with breast cancer, head and neck cancer, lung cancer, renal cancer, and cutaneous melanoma (Fig. 1a). LTβR expression levels were comparable with many oncogenes such as MYC, TP53, and BRCAs in melanoma patients (Fig. 1b), supporting that high LTβR expression on tumors denotes worse patient prognosis. Cancer patient samples were stratified as high and low LTβR expressors using an expression cut-off of one standard deviation above and below the mean of normalized gene expression. The clustering of gene expression of high and low LTβR expression cohorts of melanoma patients was assessed using principal component analysis (PCA). There was distinct clustering of samples, indicating underlying differences between LTβR high and low expressing melanomas (Fig. 1c). To identify other genes that may play an important role alongside LTβR, we analyzed differential gene expression (DGE) between the high and low LTβR expression cohorts. We identified 6,394, 6,895, and 3,651 DEGs for melanoma, breast cancer, and lung cancers, respectively, with an adjusted p-value < 0.05 and at least a 2-fold change in gene expression (Fig. 1d-f). Among all three types of cancers, we identified shared DEGs that were upregulated in LTβR high cohorts, including chemotactic chemokines (CCL20, CCL21, CXCL1, CXCL2, and CXCL5)^16^; Sox F genes (SOX7, SOX17, and SOX18) which play important roles on angio- and lymphangiogenesis^17,18^, and S100A family genes (such as S100A4 and S100A9) with well-established roles in tumor metastases and invasion^19,20^. Functional gene ontology (GO) enrichment for skin cutaneous melanoma revealed 272 GO terms with an adjusted p-value < 0.01. The top ten significantly enriched signaling activities identified in LTβR high cohorts (Fig. 1g) were related to regulation of cellular chemotaxis, immune cell activation and migration, and regulation of inflammatory responses.

**Fig. 1:**
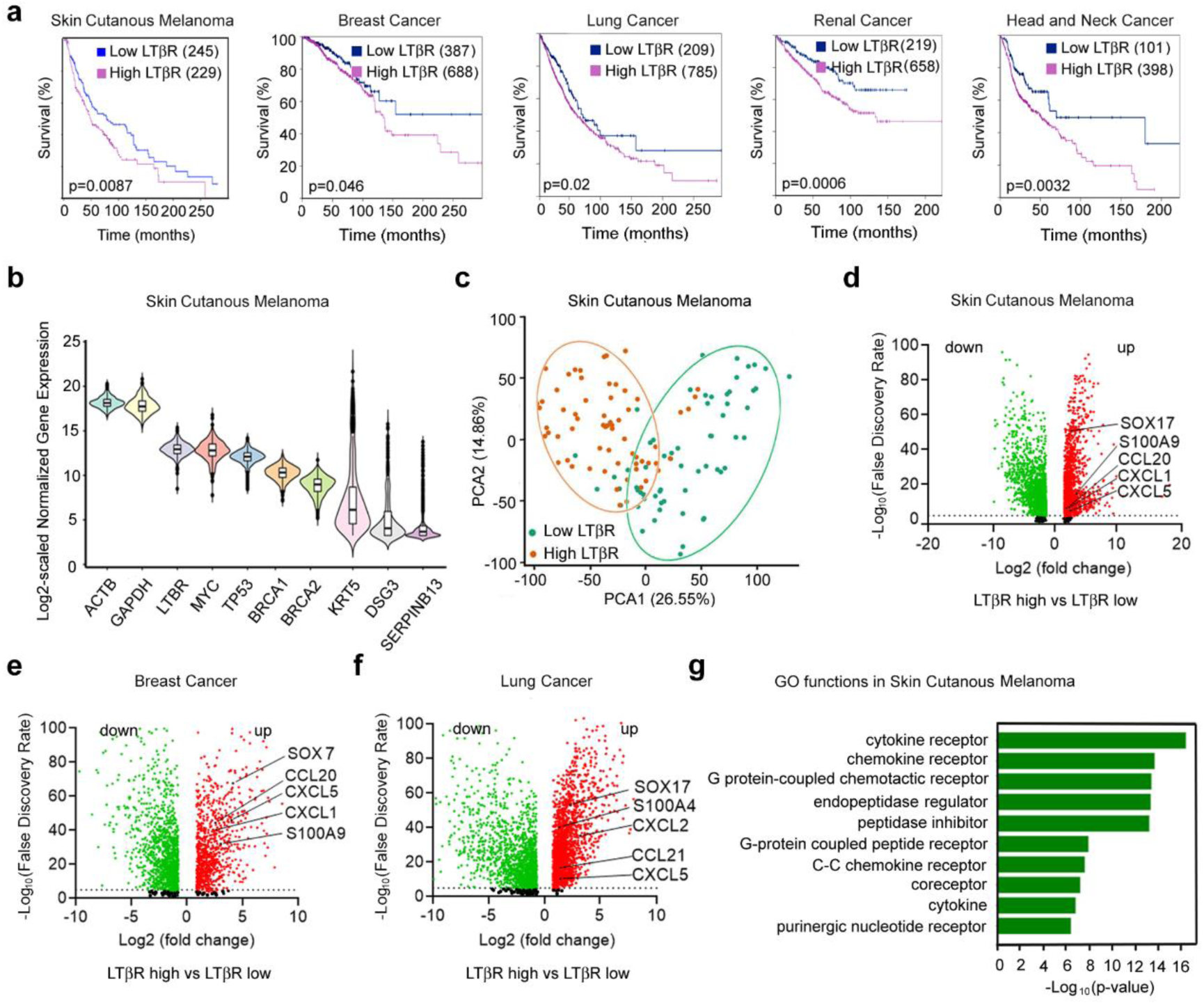
High LTβR expression in tumors is associated with poor survival in cancer patients. **a**, Kaplan-Meier survival analysis of LTβR expression (derived from Human Protein Atlas and TCGA data) in various cancer cohorts. “Low” versus “high” data segregated based on one standard deviation below or above mean expression of LTβR, Mantel-Cox log rank test. **b**, Analysis of LTβR and other candidate gene expression levels across skin cutaneous melanoma reveals that increased LTβR gene expression is associated with various oncogenes. **c**, Principal component analysis (PCA) from TCGA melanoma cohort for LTβR “high” versus “low” LTβR subsets (n = 65, 62, respectively). **d-f**, Volcano plot analysis evaluating differentially expressed genes (DEGs) from LTβR “high” versus “low” LTβR cohort of TCGA skin cutaneous melanoma (**d**), n = 6,394; breast cancer (**e**), n = 2163; and lung cancer (**f**), n = 3285, adjusted p-value < 0.05 and > 2-fold change in gene expression. **g**, Gene ontology (GO) enrichment analysis in 272 GO terms for TCGA skin cutaneous melanoma. The p-value reflect the association between a set of genes in TGCA dataset and a biological function is significant (adjusted p-value < 0.01). Top 10 statistically significant pathways are shown.

### Most tumor cells express LTβR which signals by classical and nonclassical pathways

Several murine and human tumor cells of diverse histologic origins were assessed for LTβR expression by flow cytometry. Most tumor cells expressed high levels of LTβR (Supplementary Table 1), including the well-studied B16F10 murine and human A375 melanoma cell lines (Fig. 2a). Indeed, analyzing the major component cells of human melanoma or breast cancer using published single cell RNA sequencing data, we found LTβR is highly expressed on cancer cells and endothelial cells (Extended Data Fig.1a-c). while its ligand genes LTα and LTβ are predominantly expressed on T cells and B cells, and Tregs have the highest combined expression levels of LTα and LTβ (Extended Data Fig.1d-e).

**Fig. 2:**
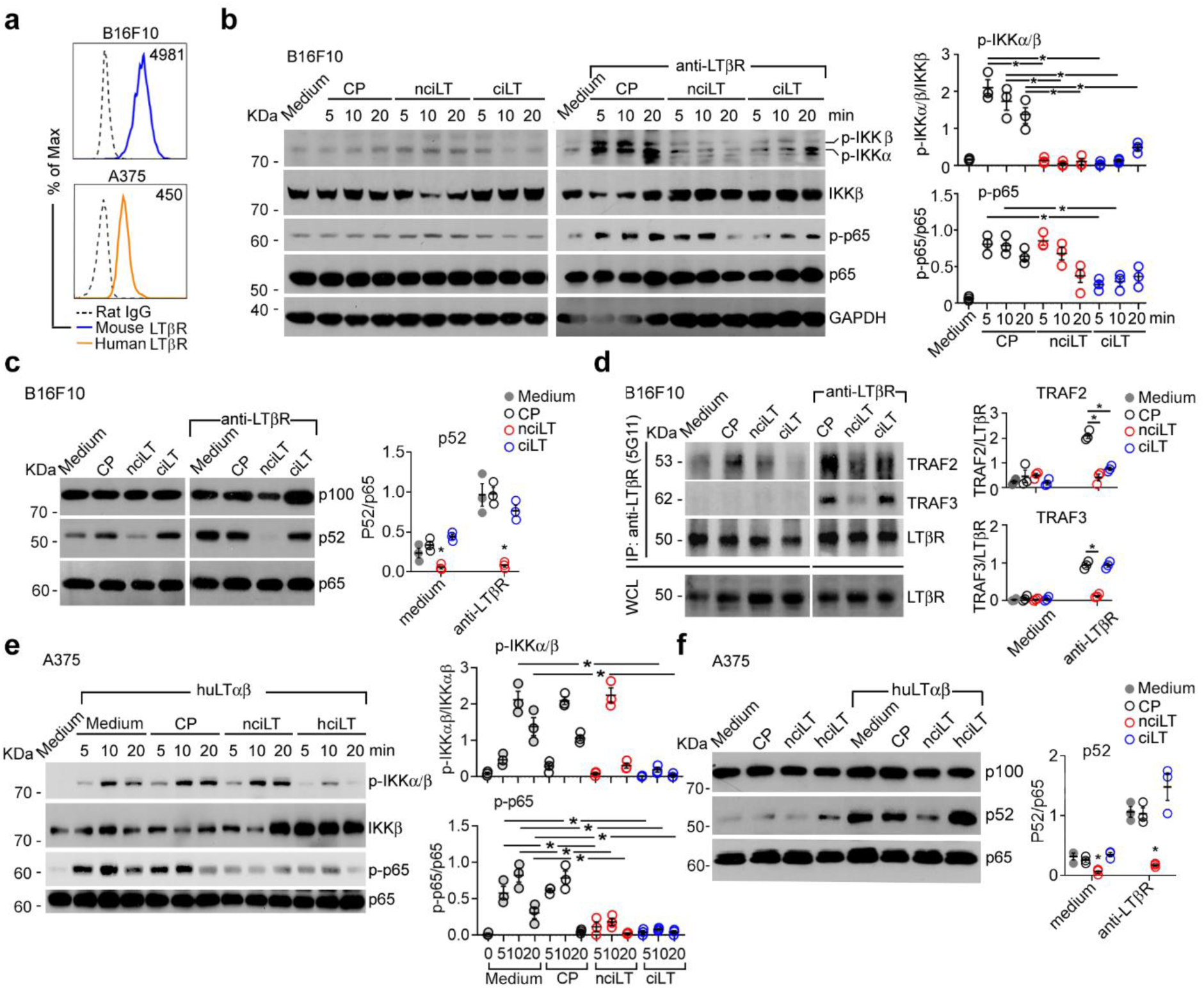
Tumor LTβR signals by classical NFκB and nonclassical NFκB pathways. **a**, Flow cytometry of LTβR expression on mouse (B16F10) and human (A375) melanoma cells. Median fluorescence intensity (MFI) shown. **b, c**, Immunoblots for classical NFκB-p65 and IKKα/β phosphorylation (**b**) or non-classical p100 processing to p52 (**c**) in B16F10 pretreated with 20 μM ciLT, nciLT, or control scrambled peptide (CP) for 1 hour at 37°C; and then stimulated with or without agonist anti-LTβR 3C8 mAb for indicated times (b) or 6 hours (**c**). **d**, Immune precipitation of B16F10 LTβR with anti-LTβR mAb (5G11). Cells stimulated with or without agonist anti-LTβR mAb for 10 minutes. Representative blots shown. WCL, whole cell lysate. **e**, **f**, Human melanoma A375 cells pretreated with 20 μM human ciLT (hciLT), nciLT, or CP for 1 hour at 37°C; and then stimulated with or without human recombinant LTαβ for indicated times (**e**) or 6 hours (**f**). Each panel representative of 3 independent experiments. **b**-**f**: Mean ± SEM. *p < 0.05, by one-way ANOVA.

To determine LTβR-mediated classical and nonclassical NFκB signaling in tumor cells, we stimulated B16F10 cells with agonist anti-LTβR mAbs along with pretreatment with cell permeable LTβR blocking decoy peptides, which we previously developed and validated to selectively block LTβR-classical NFκB-IKKβ/p65 (ciLT) or nonclassical NFκB-IKKα/NIK/p52 (nciLT) pathways^3^. Unstimulated cells in a steady state showed weak IKKα/β and NFκB-p65 phosphorylation, which were not affected by either ciLT or nciLT blocking peptides (Fig. 2b, left panel). LTβR ligation induced phosphorylation of IKKα/β (Fig. b, right panel) which was inhibited by both ciLT and nciLT 5 to 20 minutes post ligation. LTβR ligation further enhanced NFκB-p65 phosphorylation, which was inhibited as early as 5 to 10 minutes by ciLT but not nciLT (Fig. 2b, right panel). Resting B16F10 cells had constitutively processed p52 and was inhibited by nciLT only (Fig. 2c, left panel). Ligation of LTβR further enhanced p52 processing which was inhibited by nciLT but not ciLT (Fig. 2c, right panel). Immunoprecipitation of LTβR showed a low level of TRAF2 and no TRAF3 bound to the B16F10 LTβR complex in the steady state (Fig. 2d, left panel). Stimulation of LTβR recruited TRAF3, and this binding was blocked by nciLT alone (Fig. 2d, right panel). Notably, stimulation of LTβR also recruited more TRAF2 to the receptor complex which was blocked by both ciLT and nciLT (Fig. 2d, right panel). These data suggest that B16F10 LTβR nonclassical NFκB signaling involves the recruitment of both TRAF2 and TRAF3 to the LTβR complex, while LTβR-classical NFκB signaling mainly requires TRAF2 recruitment. Similar signaling patterns were observed in human melanoma A375 cells (Fig. 2e-f), Together, these data show that nciLT preferentially blocks B16F10 LTβR nonclassical NFκB signaling. Interestingly, nciLT suppressed IKKα/β but not p65 phosphorylation in mouse melanoma and suppressed p65 but not IKKα/β phosphorylation in human melanoma, indicating cellular context dependent classical NFκB signaling. It is noteworthy that the same peptide blocks TRAF recruitment to LTβR differently in tumor cells and LECs: nciLT blocks both TRAF2 and TRAF3 binding in B16F10 but blocks only TRAF3 in LECs, while ciLT blocks only TRAF2 in B16F10 but blocks both TRAF2 and TRAF3 in LECs, again indicating cellular context dependent LTβR signaling.

### Tumor LTβR nonclassical NFκB signaling preferentially regulates TEM

We previously characterized the role of LTβR in migration and its regulation by Tregs, endothelial cells^3,11,12^. Focusing on melanoma and breast cancer which follow reliable routs of lymphatic metastasis^15^, we used the specific blocking peptides to investigate which LTβR signaling pathways regulate tumor lymphatic TEM. We treated mouse and human tumor cell lines with the decoy peptides or with the agonist anti-LTβR mAb and assessed their migration across Boyden chamber transwells coated with the mouse lymphatic endothelial cell line SVEC4-10 or human primary LECs. To enhance migration, we used the chemotactic molecule sphingosine 1-phosphate (S1P), which is spatially compartmentalized in blood, lymph, and tumor, and has widely expressed receptors (S1P1-5)^21^, recapitulating migration from tissue to lymphatic vessels as would occur during metastatic spread^22^. The nonclassical LTβR-NFκB blocking peptide nciLT (identical sequence in human and mouse) but not classical LTβR-NFκB blocking peptide ciLT (versions specific for mouse and human used) was sufficient to inhibit TEM of murine tumor cells such as sarcoma KPI30, breast cancer 410, ovarian cancer ID8, mammary adenocarcinoma 66.1 (Fig. 3a), human breast cancer MDA-MB-231, and human melanoma A375 (Fig. 3a-b and Supplementary Table 2). Both nciLT and ciLT inhibited TEM of mouse melanoma B16F10 and human lung cancer A549, although nciLT was far more effective. LTβR stimulation by anti-LTβR mAb enhanced TEM of mouse or human melanoma and breast cancer (Fig. 3a-b). Notably, tumor cell TEM required the presence of LECs for migration to an S1P gradient (Fig. 3c). This is reminiscent of our previous report that T lymphocyte chemotaxis to S1P also requires the presence of LEC for efficient migration ^22^. This suggests that factors produced by LECs, such as adhesion or junctional molecules or chemokines, are critical for tumor migration.

**Fig. 3:**
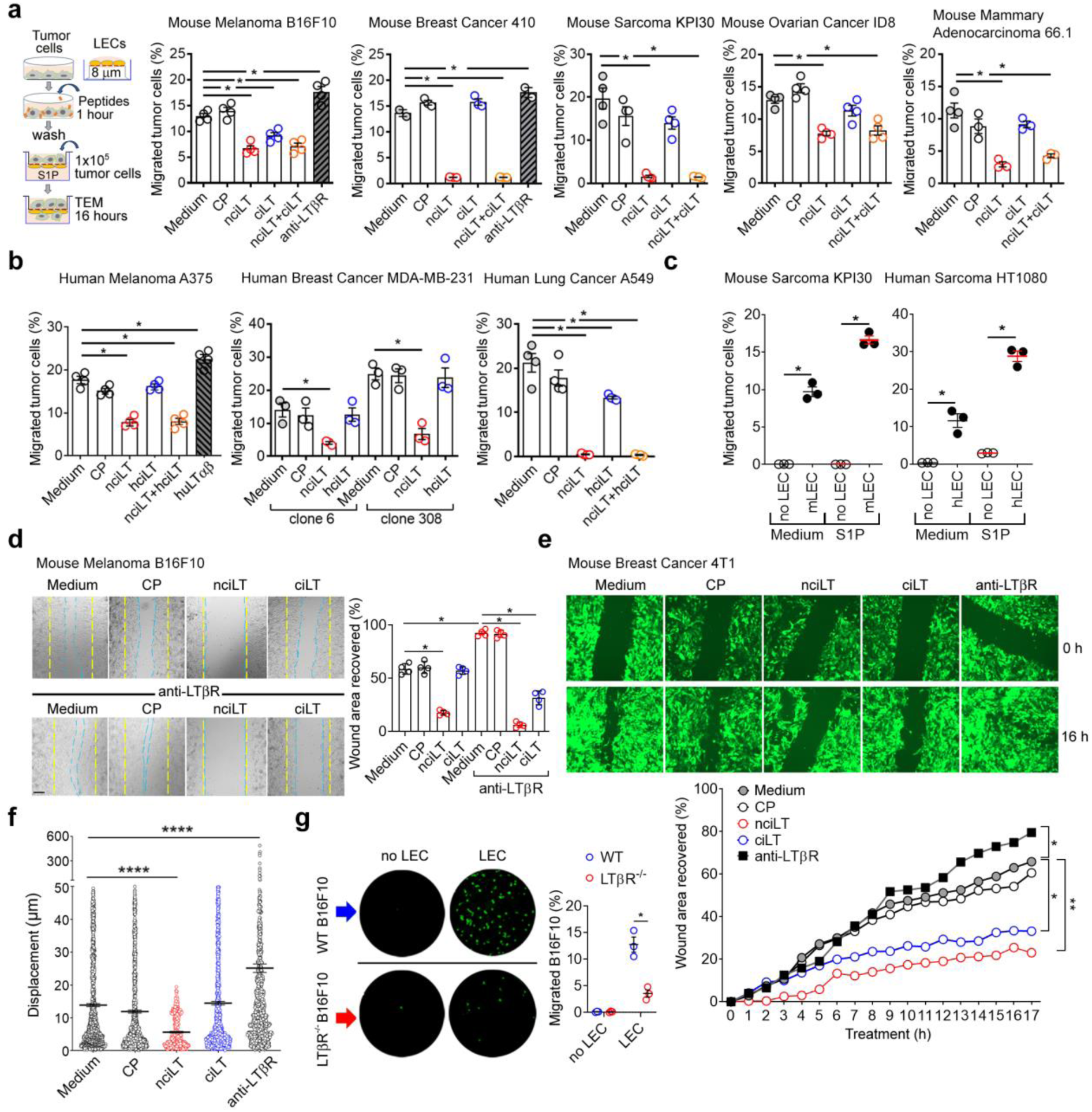
Most tumors use nonclassical NFκB signaling for lymphatic migration. **a**, **b**, Transwell TEM assay of mouse (**a**) or human (**b**) cancer cell lines. Tumor cells pretreated with 20 μM nciLT, ciLT (or hciLT), or CP for 1 hour at 37°C; or treated with 2 μg/mL agonist anti-mouse LTβR Ab (3C8) or LTβR Ig (**a**); or 10 ng/mL human LTβR ligand huLTαβ (**b**) for 1 hour at 37°C, washed and loaded for TEM across mouse or human LECs toward 200 μM S1P for 16 hours. **c**, Transwell TEM assay of mouse or human sarcoma cells toward medium or 200 μM S1P crossing the 8 μm Boyden chamber coated with or without mouse or human LECs, respectively. **d**, Images of cell migration into area of defect after scratching confluent B16F10 monolayers treated with indicated blocking peptides. Migration into area of cell defect measured after 16 hours. **e**, Representative images of mouse breast cancer 4T1-eGFP migration into area of cell defect at 0 and 16 hours after treatment with indicated peptides. Area recovery measured with the time-lapse microscopy. **f**, Time-lapse microscopy of B16F10-GFP pre-treated with peptides as in **a**-**c**. **g**, TEM of WT or LTβR^−/−^ B16F10 across transwell coated with or without mouse LECs toward 200 μM S1P for 16 hours. **a**-**e**: Representative of 3 independent experiments. Mean ± SEM. *p < 0.05, by one-way ANOVA.

### Tumor LTβR nonclassical NFκB signaling regulates cell migration and motility

Tumor cells were pretreated for 1 hour with LTβR signaling blocking peptides and assessed in a migration response assay. Consistently, blockade of B16F10 or 4T1 LTβR nonclassical NFκB signaling with nciLT abolished the migration response (Fig. 3d-e). ciLT inhibited the migration response with lesser efficiency.

Monitoring peptide-treated B16F10-eGFP migration across LECs by time-lapse microscopy, we observed that nciLT suppressed tumor cell motility assessed by cell displacement while agonist anti-LTβR mAb stimulated B16F10 motility (Fig. 3f). In addition, CRISPR/Cas9 LTβR-depleted B16F10 had impaired TEM across LECs (Fig. 3g). Overall, the data demonstrates that tumor LTβR nonclassical and to a lesser extent classical NFκB signaling plays critical roles in tumor motility and migration.

### Tumor LTβR nonclassical signaling regulates tumor growth

Since LTβR expression was associated with worse prognosis and increased expression of detrimental GO terms for tumor patients, we assessed LTβR regulation of tumor growth in vitro. Pretreatment of wild type (WT) B16F10 for 1 hour with nciLT inhibited cell growth, and inhibition was abolished in CRISPR/Cas9 LTβR depleted B16F10 (Fig. 4a), indicating that inhibition is LTβR signaling specific. Peptide treatment for 1 hour with or without LTβR stimulation by agonist mAb did not affect B16F10 cell viability (Fig. 4b). However, 5 to 16 hours of nciLT treatment increased B16F10 apoptosis (Fig. 4c), while other peptides with or without LTβR stimulation did not affect viability. Importantly, LTβR stimulation prevented the increased apoptosis by nciLT (Fig. 4c), indicating LTβR stimulation induced classical NFkB signaling promoted tumor cell survival and compensated for nciLT triggered apoptosis. These data indicate nciLT inhibits tumor cell growth and promotes apoptosis.

**Fig. 4:**
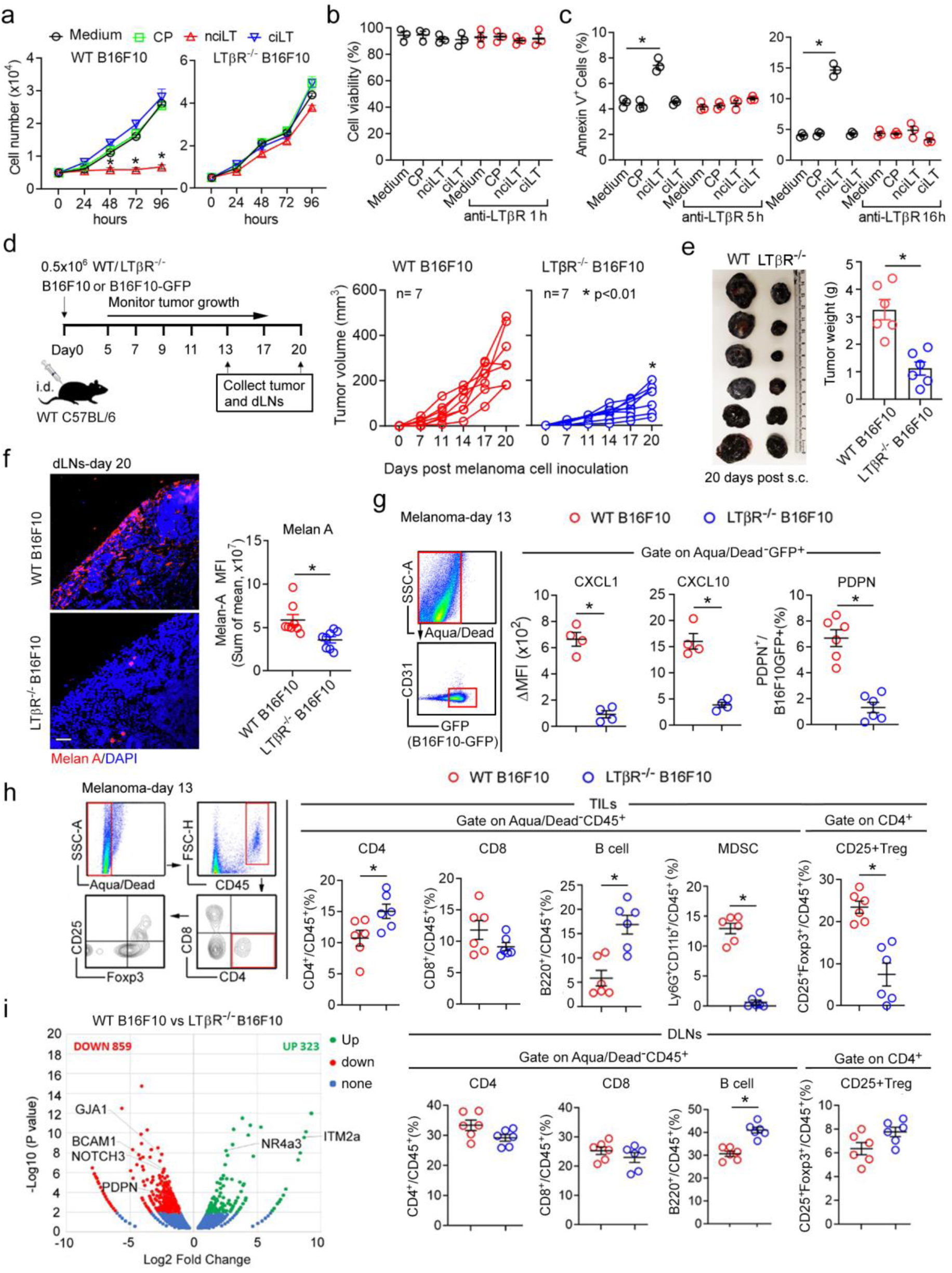
Tumors use LTβR nonclassical NFκB signaling for tumor growth. **a**, In vitro cell growth of WT or LTβR^−/−^B16F10 pretreated with 20 μM of CP, nciLT, or ciLT for 1 hour. **b**, Cell viability analysis of B16F10 treated with indicated peptide with or without anti-LTβR mAb for 1 hour and then assessed by MTT assay. **c**, Flow cytometry apoptotic cell analysis of Annexin V^+^ B16F10 pretreated with indicated peptides for 1 hour, and then stimulated with or without anti-LTβR mAb for 5 or 16 hours. **d**-**h**, C57BL/6 mice intradermally transferred with WT or CRISPR/Cas9 LTβRKO (LTβR^−/−^) B16F10. Scheme of experimental setup and tumor growth (**d and e**). At day 20, dLNs analyzed for Melan-A expression of metastatic B16F10 cells in dLNs by immunohistochemistry (**f**). Magnification 20x; scale bar 42 μm. At day 13, transferred B16F10GFP tumors analyzed for CXCL1, CXCL10, and PDPN expression in GFP^+^B16F10 cells (**g**), and dLNs analyzed for CD4, CD8, Foxp3^+^ CD4 Tregs, B220^+^B cells, CD11b^+^Ly6G^+^MDSCs (**h**). **i**. Volcano plot of differential gene expression comparing LTβR^−/−^ to WT B16F10. Genes upregulated (green) or down regulated (red) at log2 fold change ≥2 or ≤−2 and P value < 0.05 adjusted by the Benjamin and Hochberg method. Three (**a**-**c**) or two (**d**-**h**) independent experiments. **a**-**h**: Mean ± SEM. *p < 0.05; **p < 0.01, by one-way ANOVA.

### Tumor LTβR depletion suppresses tumor growth and metastasis

CRISPR/Cas9 LTβR-depleted B16F10 cells were assessed for tumor growth and migration. Subcutaneously inoculated LTβR-depleted B16F10 cells had slower in vivo growth than WT B16F10 (Fig. 4d-e) and reduced lymphatic metastasis into the draining LNs (dLNs) (Fig. 4f). LTβR-depleted B16F10 melanoma had impaired production of tumor cell-derived CXCL1 and CXCL10 (Fig. 4g), and reduced intratumoral Tregs and MDSCs but increased CD4 T cells, B cells, and unaltered CD8 T cells 13 days after melanoma inoculation, compared to WT B16F10 (Fig. 4h). Since these cells were not decreased in tumor dLNs, this suggests that LTβR depletion in tumor cells had no impact on immune cell egress from the tumor. However, the reduced Tregs and MDSCs may result from the reduced CXCL1 and CXCL10 which may recruitment these CXCR2 and CXCR3 expressing cells to tumor from the blood circulation (Fig. 4h).

We explored the genes affected by LTβR depletion by whole transcriptome RNA sequencing (RNA-seq) analysis, comparing CRISPR/Cas9 LTβR^−/−^ to WT B16F10. DEGs were observed between the two cell lines with 323 upregulated and 859 downregulated in LTβR^−/−^ B16F10 (Fig. 4i). The enhanced expression of tumor suppressor genes such as integral membrane protein 2A (*ITM2a*) and nuclear receptor 4A3 (*NR4a3*) in LTβR-depleted B16F10 cells may contribute to cell growth inhibition, while the decreased gap junction protein alpha 1 (*GJA1*), podoplanin (*PDPN*), *NOTCH3*, and basal cell adhesion molecule 1 (*BCAM1*) which regulate cell adhesion may contribute to reduced Treg tumor recruitment and melanoma metastasis. In addition, GO analysis indicated predominant downregulation of cell surface G-protein-coupled receptor (GPCR) signaling pathways that regulate chemotaxis, localization, and transmembrane transport, in addition to cellular component organization or biogenesis and developmental regulations as key changes induced by LTβR depletion (Extended Data Fig.2a-c).

### Genes driven by tumor LTβR NFκB signaling

To determine the molecules regulated by nonclassical LTβR-NFκB signaling in B16F10 melanoma, we employed bulk RNASeq in a two-pronged strategy. First, we performed RNA-Seq on B16F10 cells stimulated by agonist anti-LTβR Ab (3C8) for 6 hours to induce LTβR nonclassical NFκB signaling. These stimulated cells were compared to both non-stimulated B16F10 cells and to cells that were stimulated and pretreated with the LTβR-nonclassical blocking peptide nciLT. Second, we generated CRISPR/Cas9 knockout (KO) of nonclassical NFκB/NIK B16F10 and stimulated them for 6 hours. Thus, we were able to compare gene expression regulated by the LTβR nonclassical NFκB pathway using both the nciLT inhibitor and NIK deficiency. Melanoma LTβR nonclassical NFκB signaling upregulated angiogenic and immunosuppressive myeloid chemokines CXCL1^20,23^, CXCL10^24^, and CCL5^25^, which were down regulated by nciLT and diminished in NIK-deficient B16F10 cells (Fig. 5a-b). In addition, protumor interferon-stimulated genes ISG15^26^ and IFIT3^27^ (interferon induced protein with tetratricopeptide repeats 3), and oncoprotein USP18^28^ (Ubiquitin Specific Peptidase 18), which suppresses IFN responses, were also driven by LTβR-nonclassical signaling, down regulated by nciLT, and diminished in NIK-deficient B16F10 cells (Fig. 5a). The selected genes were further confirmed by real-time PCR (Fig. 5b). CXCL1, CXCL10 and CCL5 expression were abolished in NIK-depleted but not IKKβ-depleted B16F10 (Extended Data Fig. 3a-b), confirming that these genes are regulated by the LTβR-non-classical NFκB NIK signaling pathway. CXCL1 and CXCL10 expression were enhanced in Treg cocultured human melanoma A375 and human LECs and inhibited by nciLT pretreatment (Extended Data Fig. 3c-d). Neither Teff coculture nor ciLT pretreatment affected the chemokine expression. Together, the data showed CXCL1 and CXCL10 production in tumor cells and LECs are driven by the LTβR-non-classical NFκB NIK signaling pathway.

**Fig. 5:**
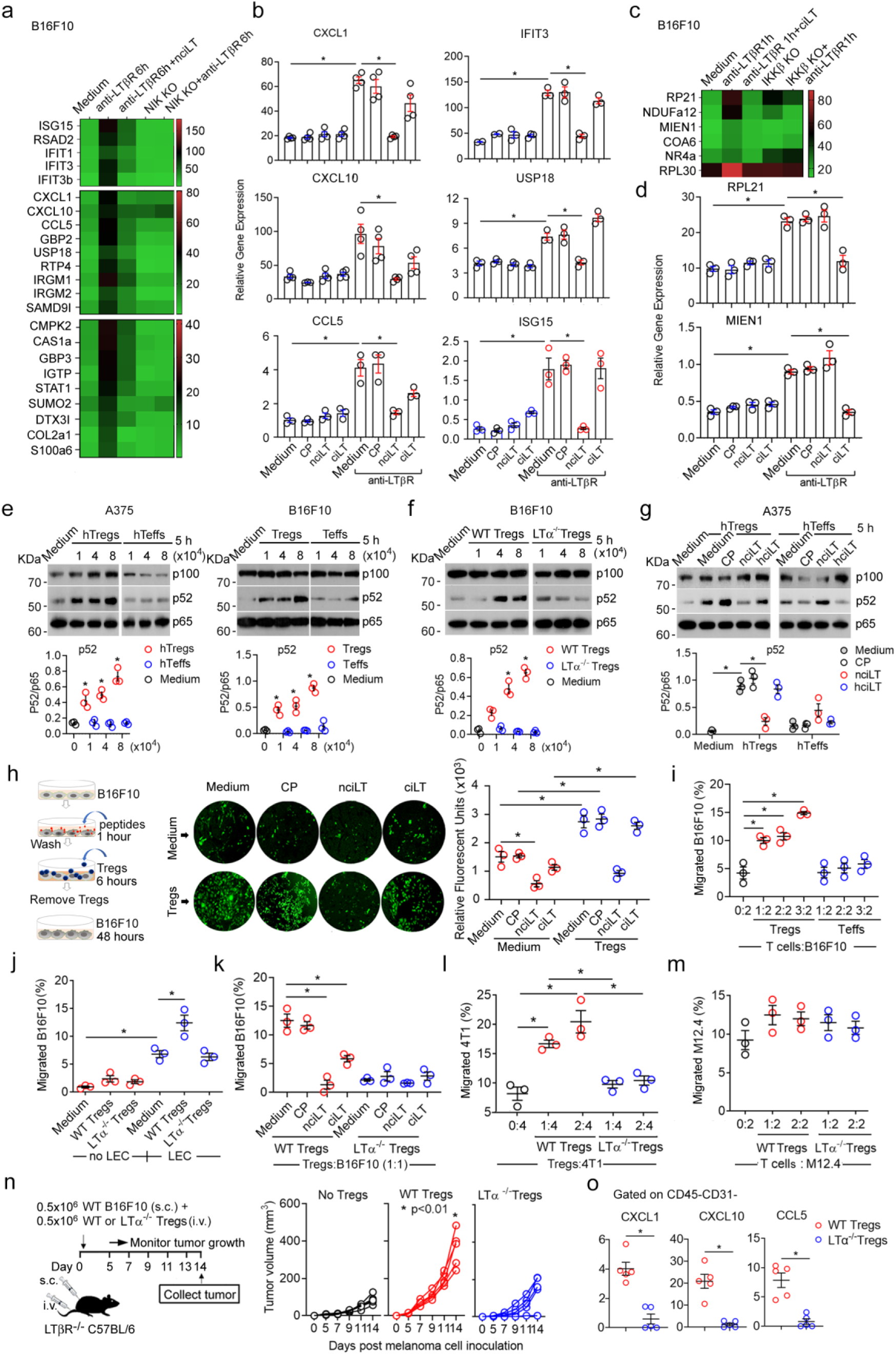
Treg LTα1β2 stimulates tumor LTβR nonclassical NFκB pathway to promote tumor growth and TEM. **a**-**d,** Bulk RNA-Seq analysis of B16F10. Genes down regulated by nciLT (**a**) or ciLT (**c**) and regulated by NIK (**a**) or IKKβ (**c**)-deficiency. WT or CRISPR/Cas9 NIK KO B16F10 pretreated with nciLT for 1 hour followed by 6-hour anti-LTβR (3C8) stimulation (**a**, **b**). WT or CRISPR/Cas9 IKKβ KO B16F10 treated with ciLT for 1 hour prior to 1 hour LTβR stimulation (**c**, **d**). Selected genes confirmed by RT-PCR for nonclassical (**b**) and classical (**d**) regulation. **e**-**g**, Immunoblots for p100/p52 in human A357 (**e**, **g**) and mouse B16F10 (**e**, **f**) melanoma cells cocultured with various dose of human or mouse Tregs, Teffs, or LTα-deficient (LTα^−/−^) Tregs as indicated for 5 hours. A375 pretreated with 20 μM CP, nciLT, or hciLT for 1 hour prior to coculturing with human T cells (**g**). **h**, B16F10 pretreated with 20 μM CP, nciLT, or ciLT for 1 hour, washed, and cocultured with Tregs for 6 hours, washed and grown for 48 hours. **i**-**m,** Migration of B16F10 cocultured with Tregs or Teffs (**i**), or with LTα^−/−^ Tregs (**j and k**) as indicated for 6 hours, washed, and migrated across LEC toward 200 μM S1P for 16 hours. B16F10 pretreated with indicated peptides before coculturing with Tregs and TEM as in h (**k**). TEM of breast cancer 4T1 cells (**i**) or B cell lymphoma M12.4 cells (**m**) cocultured with WT or LTα^−/−^ Tregs. **n-o**, 0.5×10^6^ WT or LTα-deficient (LTα^−/−^) Tregs intravenously transferred to LTβRKO (LTβR^−/−^) C57BL/6 mice which were subcutaneously inoculated with 0.5×10^6^ WT B16F10. Scheme of experimental setup and tumor growth (**n**). At day 14, tumors analyzed for CXCL1, CXCL10, and CCL5 expression. b, d, e-m, and o, data representative of 3 independent experiments. Mean ± SEM. *p < 0.05 by one-way ANOVA.

A parallel strategy was employed for identifying LTβR-classical NFκB-driven genes, by comparing gene expression in cells stimulated for one hour with agonist mAb and pretreated or not with the classical blocking peptide ciLT, and further compared with IKKβ-depleted B16F10 (Fig. 5c). Several pro-cancer genes such as ribosomal proteins 21 and 30 (RPL21 and RPL30)^29^, migration and invasion enhancer 1 (MIEN1)^30^, and nuclear receptor NR4a1^31^ were identified to be rapidly induced by LTβR activation and inhibited by blocking LTβR classical NFκB signaling (Fig. 5c). Selected genes RPL21 and Mien1 were further shown by real-time PCR to be inhibited by the classical NFκB blocking peptide ciLT but not by the nonclassical NFκB blocking peptide nciLT (Fig. 5d), confirming regulation by the LTβR-classical NFκB signaling pathway.

### Treg LTα1β2 stimulates tumor LTβR nonclassical NFκB signaling to enhance tumor growth

Our previous study revealed that Tregs express high levels of LTα1β2 which ligate and activate LTβR on LEC^12^. Treg LTα1β2 signals primarily to the LTβR nonclassical NFκB pathway and alters LEC morphology and membrane characteristics, thereby causing endothelial morphologic changes and promoting lymphatic TEM of other immune cells ^12^. To explore the possible role of Tregs directly interacting with tumor cells, we assessed whether Treg LTα1β2 can ligate and activate tumor surface LTβR. LTα1β2 high expressing human Tregs and LTα1β2 low expressing human effector T cells (Teffs) (Extended Data Fig. 4a) cocultured with the melanoma cells equally induced phosphorylation of IKKα/β and p65 classical NFκB signaling in human A375 cells as early as 10 minutes after incubation (Extended Data Fig.4b). An identical pattern was observed in mouse B16F10 cultured with mouse Tregs and Teffs (Extended Data Fig. 4c). After 5 hours coculture with both human and mouse melanoma cells, only Tregs induced non-classical NIK signaling for p100 processing to p52 in these tumor cells, while LTαβ-low expressing Teffs (Fig. 5e) or LTα-deficient Tregs (Fig. 5f) did not. Only nonclassical blocking peptide inhibited Treg induced nonclassical NFκB signaling in human melanoma(Fig. 5g), while both nciLT and ciLT inhibited the Treg induced nonclassical signaling in B16F10 (Extended Data Fig. 4d). Other immune cells such as B cells, CD8 T cells also express LTα1β2, however, the expression level is significantly lower than that of Tregs (Extended Data Fig. 4e), and B cells failed to induce any NFkB signaling in either tumor or LECs (Extended Data Fig. 4f) and had no impact on tumor cell TEM (Extended Data Fig.4g). These results indicate that Treg LTαβ exclusively induces nonclassical NFκB signaling on tumor cells through cell-cell interactions. To investigate the biological function of Treg-induced preferential LTβR nonclassical NFκB signaling, we cocultured Tregs with B16F10 which were pretreated with or without blocking peptides to assess tumor cell growth. Treg-cocultured B16F10 demonstrated increased cell growth. Blockade of only the nonclassical NFκB signaling abolished Treg stimulated growth and likewise inhibited basal growth of tumor cell even without Treg pretreatment (Fig. 5h), similar to the observations in Fig. 4a. Thus, these data show that both basal constitutive and Treg induced LTβR-nonclassical-NFκB signaling in B16F10 promote tumor growth.

### Treg LTα1β2 stimulates tumor LTβR nonclassical NFκB signaling to enhance tumor cell TEM

To determine whether Tregs directly affect tumor cell migration, we cocultured Tregs or Teffs with melanoma cells prior to migration assay across LEC coated transwell. Treg-cocultured B16F10 cells showed enhanced TEM across LECs in a dose-dependent fashion, while Teffs induced no such effect (Fig. 5i). Human Tregs co-cultured A375 also had increased TEM (Extended Data Fig. 4h). Importantly, the effect required the presence of LECs. Without LECs, the Treg-pretreated B16F10 had no enhanced TEM (Fig. 5j). Treg enhanced TEM was LTα1β2-LTβR signaling dependent since LTα-deficient Tregs failed to increase tumor TEM (Fig. 5k). Blockade of B16F10 LTβR nonclassical NFκB signaling also abolished Treg promoted tumor TEM (Fig. 5k). Similarly, Tregs but not LTα-deficient Tregs, enhanced 4T1 breast cancer TEM (Fig. 5l). Treg failed to increase TEM of B cell lymphoma M12.4 which has minimal levels of LTβR expression (Fig. 5m and Supplementary Table 1). Together, these data show that Treg LTαβ directly ligates tumor LTβR to activate nonclassical NFκB signaling and enhance tumor cell TEM. To assess in vivo Tregs direct interaction with tumor cells, we intravenously injected WT or LTα-deficient Tregs to LTβR-depleted mice inoculated with WT B16F10. Thus, in this system, Treg LTα1β2 can only ligate and activate tumor surface LTβR. WT Tregs significantly promoted melanoma growth, while LTα-deficient Tregs failed to do so (Figure 5n). There was also increased LTβR nonclassical signaling driven CXCL1, CXCL10, and CCL5 expression in melanoma cells in the host mice injected with WT Tregs but not with LTα-deficient Tregs (Figure 5o). These results demonstrate that in vivo Treg LTα1β2 can directly activate tumor cell LTβR nonclassical NFκB signaling.

### Treg LTα1β2 stimulates LEC LTβR nonclassical NFκB to promote tumor growth and TEM

We previously demonstrated that Tregs stimulate LTβR nonclassical NFκB signaling in LECs and thereby alter LEC characteristics for leukocyte TEM^12^. Since we showed above that Tregs also stimulate tumor LTβR to alter tumor growth and TEM, we next analyzed if this signaling induced additional events between LECs and tumors. LECs were pretreated with Tregs, the Tregs removed and then B16F10 added to the LECs. This coculture promoted melanoma growth (Fig. 6a). Blockade of LEC LTβR nonclassical NFκB with nciLT prior to coculture with Tregs abolished the activated LEC enhanced tumor cell growth (Fig. 6a, middle row). Blockade of LEC LTβR-classical NFκB with blocking peptide ciLT had no effect on tumor cell growth. LECs pretreated with Teffs did not affect tumor cell growth, and the blocking peptides did not alter the lack of effect of Teffs on LEC for tumor growth (Fig. 6a, lower row). To determine whether the growth effect was from direct tumor-LEC contact or due to the secreted factors from the LECs, 0.4 μm pore transwells were used to separate B16F10 in the upper chamber from LECs in the lower chamber. The B16F10 or the LECs were each independently cocultured with Tregs for 6 hours, the Tregs then removed, and the growth of B16F10 assessed after another 48 hours. WT or LTα-deficient Tregs cocultured with LECs in the lower chamber did not promote B16F10 cell growth in the upper chamber (Extended Data Fig. 5a-b), suggesting soluble cytokines/chemokines produced by activated LECs have no effect on B16F10 cell growth, and that a direct tumor-LEC cell physical contact is required for supporting tumor cell growth. To test the effect of Treg LT-triggered LEC LTβR signaling on tumor migration, we co-cultured Tregs with WT or LTβR-deficient LEC prior to assessing tumor cell TEM. WT LECs co-cultured with Tregs promoted B16F10 TEM, while LTβR-deficient LECs co-cultured with WT Tregs did not enhance B16F10 migration (Fig. 6b). WT LECs co-cultured with LTα-deficient Tregs also had no effect on B16F10 or 4T1 TEM (Fig. 6c).

**Fig. 6:**
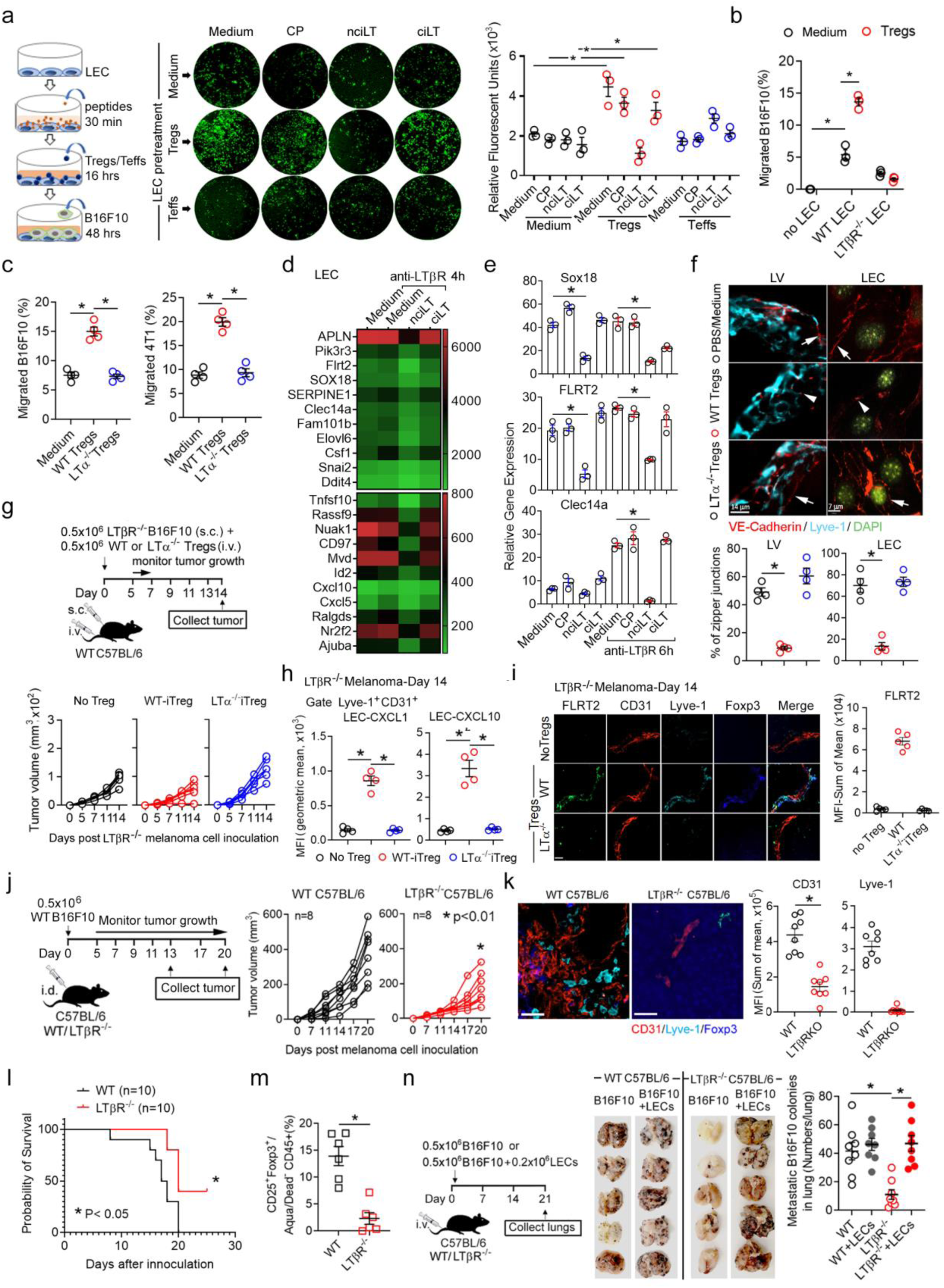
Tregs communicate with LEC through LTα1β2/LTβR signaling to activate nonclassical NFκB pathway and promote tumor growth and TEM. **a**, Peptide-pretreated LEC cocultured with Tregs or Teffs (LEC: T cell = 1: 2) for 16 hours, followed by removal of T cells, and then LEC cocultured with B16F10-GFPs (LEC: B16F10 = 1:1) for 48 hours. B16F10-GFPs analyzed by EVOS microscopy. **b, c,** Transwell assay, B16F10-GFPs migrated across WT or LTβR^−/−^ LECs pre-treated with or without Tregs as in **a** (**b**). B16F10 or 4T1 cells migrated across LECs pretreated with WT or LTα^−/−^Tregs for 16 hours (**c**). **d,** Heatmap of gene array analysis of mouse primary dermal LECs treated with nciLT and ciLT. LECs treated with 20 μM nciLT or ciLT for 30 minutes prior to LTβR stimulation with anti-LTβR Ab (3C8) for 4 hours. **e**, Regulated genes selected for real-time PCRs. **f,** Immunohistochemistry of intercellular VE-cadherin in lymphatic vessels (LV) or cultured LECs pretreated with WT or LTα^−/−^Tregs for 16 hours. Magnification 60x; scale bar:14 μm (LV) and 7 μm (LEC). **g-i**, Intravenous transfer of WT or LTα^−/−^Tregs (5×10^5^) to WT C57BL/6 mice subcutaneously inoculated with LTβR KO B16F10 (5×10^5^). Scheme of experimental setup and tumor growth (**g**). At day 14, tumors analyzed for CXCL1, CXCL10 expression in LECs by flow cytometry analysis (**h**); FLRT2 expression in LECs by Immunohistochemistry (**i**). **j-m**, Intradermal transfer of B16F10 cells (5×10^5^) to WT or LTβR^−/−^ C57BL6 mice. Tumor growth (**j**), tumor blood (CD31^+^Lyve-1^−^) and lymphatic (Lyve-1^+^) vessels by immunohistochemistry (**k**), mouse survival (**l**), and intratumoral Foxp3^+^CD25^+^Tregs by flow cytometry (**m**). Magnification 20x; scale bar: 40 μm. **n**, Lung colonization assessed at 3 weeks after tail vein injection of 3×10^5^ B16F10 cells with or without primary mouse LECs (1×10^5^) to WT or LTβR^−/−^ C57BL6 mice. **a-c, f,** representative of 3 (a-c, f,) and 2 (g, j, n) independent experiments shown. Mean ± SEM. *p < 0.05 by one-way ANOVA.

### Genes driven by LTβR nonclassical NFκB signaling in LEC

To investigate which molecules driven by LEC LTβR signaling to regulate tumor cell growth and migration, we employed DNA microarray analysis of LECs. Primary LECs were treated with control peptide (CP), nciLT, or ciLT prior to agonist anti-LTβR mAb stimulation. Endothelial specific genes which promote tumor malignancy and metastasis, such as fibronectin leucine rich transmembrane protein 2 (FLRT2), SRY-box transcription factor 18 (SOX18), and C-type lectin domain containing 14A (Clec14a), were identified to be driven by LTβR nonclassical NFκB signaling and were down regulated by nciLT but not by ciLT (Fig. 6d). The selected genes were further confirmed by RT-PCR (Fig. 6e). Some genes were not transcriptionally regulated while their functions relied on post-translational changes, such as VE-cadherin which is important for endothelial integrity^32^. Thus, 16 hours pretreatment with Tregs could transform intact zipper junctional VE-cadherin expression in lymphatic vessels (LVs) *in vivo* or LECs *in vitro* to button-like junction, while LT-deficient Tregs failed to do so (Fig. 6f).

### Treg LTα1β2 activates LTβR nonclassical NFkB signaling on tumor associated LECs *in vivo*

To study the direct in vivo influence of Treg LTα1β2 on LECs, we intravenously transferred 5 ×10^5^ WT or LTα^−/−^ Tregs to WT C57BL/6 mice which were subcutaneously inoculated with 0.2×10^6^ LTβRKO B16F10. In this system, Tregs can only ligate and activate surface LTβR on stromal cells, including LECs, but not on tumor cells. Since the LTβR deficient B16F10 had impaired tumor growth compared to WT B16F10 (Fig. 4d), WT Tregs did not significantly promote LTβR deficient melanoma growth (Fig. 6g), although it promoted WT B16F10 (Fig. 5n). However, WT Treg but not LTα^−/−^ Treg transfer induced more CXCL1 and CXCL10 in the tumor LECs (Fig. 6h). WT Tregs but not LTα^−/−^ Tregs accumulated around tumor lymphatic vessels where the nonclassical LTβR-NIK signaling driven endothelial specific proteins FLRT2, SOX18, and Clec14 were highly upregulated (Fig. 6i and Extended Data Fig. 6a). These data indicated that in vivo Treg LTα1β2 can directly activate LTβR nonclassical NFκB signaling in tumor LECs. To study the combined in vivo influence of Treg LTα1β2 on tumor cells and LECs for tumor growth, we intravenously transferred 1×10^6^ WT or LTα^−/−^ Tregs to WT C57BL/6 mice transplanted with 0.5×10^6^ B16F10. WT Tregs significantly promoted melanoma growth, while LTα^−/−^ Tregs failed to do so, suggesting the stronger impact of Treg LTα1β2 on tumor cells than on LECs for LTβR-mediated tumor growth (Extended Data Fig.6c).

To study whether in vivo LTβR depletion affected tumor growth, we transplanted WT B16F10 into WT or LTβRKO C57BL6 mice, which have lymphatics but lack LNs^33^. Transplanted tumors in LTβRKO mice had significantly reduced tumor growth, angiogenesis, and lymphangiogenesis compared to those in WT mice (Fig. 6j-k), and survival was also significantly higher in LTβR-deficient mice (Fig. 6l). Analysis of TILs in LTβR KO mice revealed reduced CD25^+^Foxp3^+^Tregs (Fig. 6m), Ly6G^+^CD11b^+^MDSCs, and F4/80^+^CD11b^+^macrophages, and increased total CD4 T cells, IFNγ^+^CD8 T cells, and B220^+^B cells compared to those of WT mice, although total CD45^+^ immune cells and CD8 T cells remain unchanged (Extended Data Fig.6b). These data suggest that recipient LTβR regulated recruitment and/or retention of Tregs, MDSCs, and macrophages into the tumor. Since LTβR-deficient mice lack LNs, tumor lymphatic metastasis could not be directly observed. ^33^ Therefore, we then injected B16F10 i.v. to study tumor pulmonary metastasis. LTβR-deficient mice had significantly reduced metastatic colonies in lung compared to WT mice. Importantly, when LTβR-deficient mice were injected i.v. with B16F10 together with WT LTβR-sufficient primary mouse LECs, there were remarkably increased pulmonary metastases (Fig. 6n), suggesting that LTβR on LECs plays critical roles for tumor metastases.

### Selective blockade of LTβR classical and non-classical NFκB inhibits tumor growth and lymphatic metastasis

LTβR signaling was assessed for its effects on tumor growth and metastasis by using the blocking peptides in mouse melanoma (Fig. 7a). nciLT dramatically suppressed melanoma growth, while ciLT treated melanoma had early regression but relapsed later (Fig. 7b). nciLT or ciLT treated melanoma at day 13 after B16F10 intradermal inoculation had increased intratumoral CD4 and CD8 T cell infiltration (Fig. 7c, upper panel). In addition, both nciLT and ciLT treated tumors had increased IFNγ-producing CD8 T cells, but no significant changes in IFNγ^+^ CD4 T cells (Fig. 7c, middle panel). nciLT but not ciLT significantly reduced intratumoral activated CD25^+^ Tregs (Fig. 7c, lower panel). In contrast, ciLT increased the CD25^−^ Treg subset. Both peptides significantly reduced granulocytic Ly6G^+^CD11b^+^ MDSC and B220^+^ B cell infiltration at day 13. Notably, only nciLT suppressed melanoma and endothelium derived CXCL1 and CXCL10 expression (Fig. 7d). Moreover, both blocking peptides suppressed tumor angiogenesis and lymphangiogenesis by reducing CD31^+^lyve-1^−^ blood vessels and CD31^+^lyve-1^+^ lymphatic vessels (Fig. 7e-g). Importantly, LTβR-nonclassical NFκB-driven SOX18, a crucial transcriptional regulator of angiogenesis ^34^, and FLRT2 which are highly expressed in LECs and facilitate tumor aggressiveness ^35,36^, were significantly reduced in lymphatic vessels in nciLT treated melanoma. ciLT also suppressed their expression but less effectively than nciLT (Fig. 7e-g). SOX18 localized in both blood (CD31^+^Lyve-1^−^) and lymphatic vessels (CD31^+^Lyve-1^+^), while FLRT2 mainly localized on lymphatic vessels alone (Fig. 7e-f, 7h). In dLNs, nciLT reduced both CD4 and CD8 T cells, while ciLT reduced CD4 but not CD8 T cells (Fig. 8a, upper panel), suggesting nciLT inhibited CD4 and CD8 T cell tumor egress, while ciLT inhibited CD4 T cell tumor egress which may contributed to the increased intratumoral CD4 Tregs. Total number of Tregs in dLNs were significantly decreased by both nciLT and ciLT (Fig. 8a, lower panel; 8b). Analysis of dLNs 20 days after melanoma development revealed that both nciLT and ciLT treated mice had significantly reduced dLN melanoma metastasis and Tregs (Fig. 8b). The increased retention of IFNγ-producing CD8 T cells in tumor-infiltrating lymphocytes (TILs) and reduced CD25^+^Tregs in tumor and dLNs in LTβR blocking peptides treated melanoma may play critical roles for anti-tumor immunity.

**Fig. 7:**
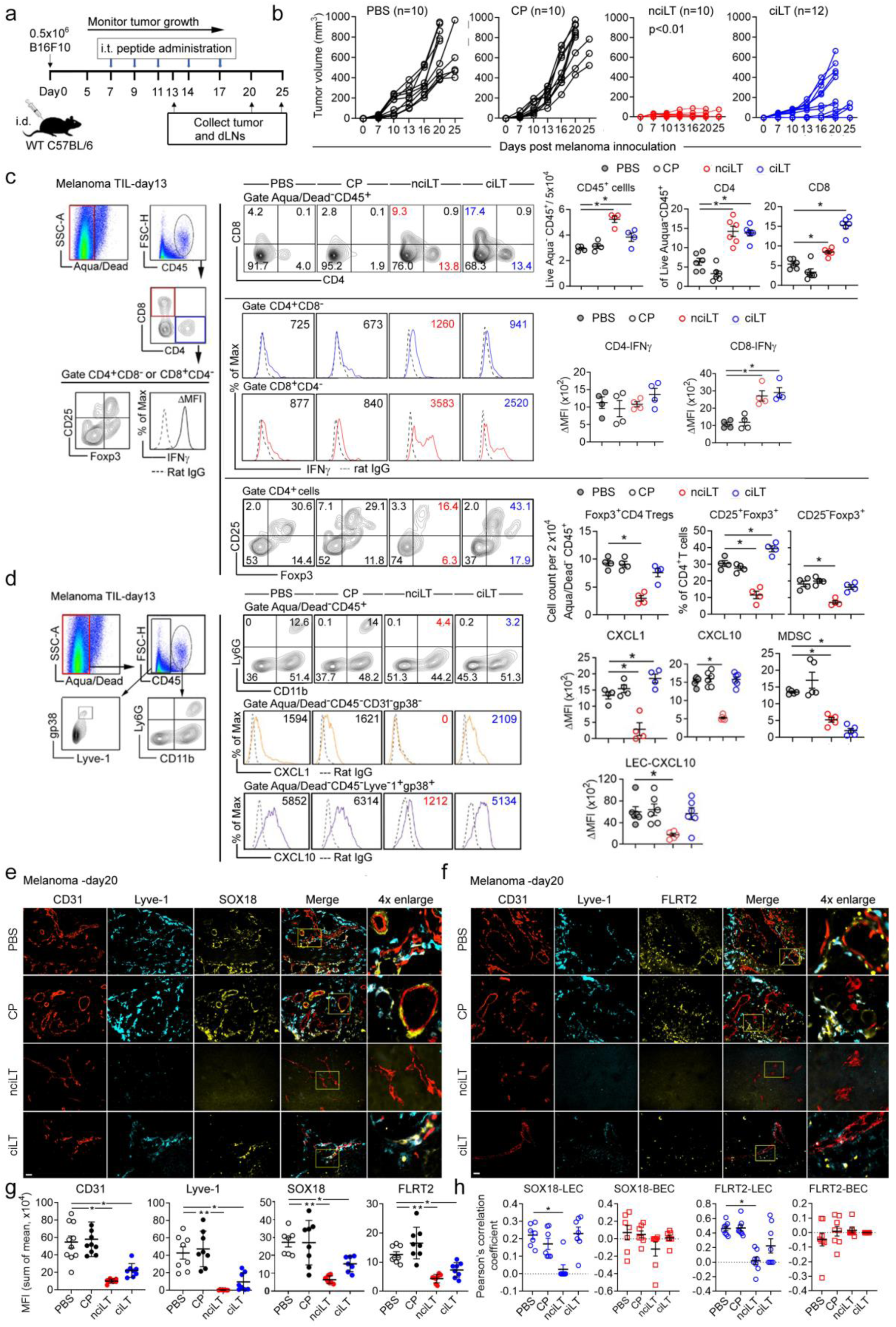
Blockade of LTβR non-classical NFκB inhibits tumor growth, immune suppressive cell recruitment and tumor angio- and lymphangiogenesis. **a, b,** Intradermally transferred B16F10 melanoma in C57BL6 mice treated with intratumoral injection of 10 nmol/tumor of nciLT, ciLT, or CP for 5 days. Scheme of tumor treatment (**a**). Tumor growth (**b**). **c, d**, At day 13, tumors analyzed for CD4, CD8, Foxp3^+^ CD4 Tregs and T cell IFNγ (**c**), Ly6G^+^CD11b^+^ MDSCs, CXCL1, and CXCL10 expression in CD45^−^ CD31^−^gp38^−^ cells or Lyve-1^+^ gp38^+^ LECs (**d**) by flow cytometry. **e**, **f,** At day 20, tumors assessed for angiogenesis (CD31^+^/Lyve-1^−^) and lymphangiogenesis (CD31^+^/Lyve-1^+^) by immunohistochemistry and analyzed for SOX18 (**e**) and FLRT2 (**f**). Magnification 20x, scale bar:42 μm. **g**, **h,** Expression on CD31^+^/Lyve-1^+^ lymphatic vessels and CD31^+^/Lyve-1^−^ blood vessels of tumors. MFIs (**g**) and quantification of co-localization between SOX18 or FLRT2 and lymphatic or blood vessels (**h**). **c**-**h**: Mean ± SEM. *p < 0.05 by one-way ANOVA.

**Fig. 8:**
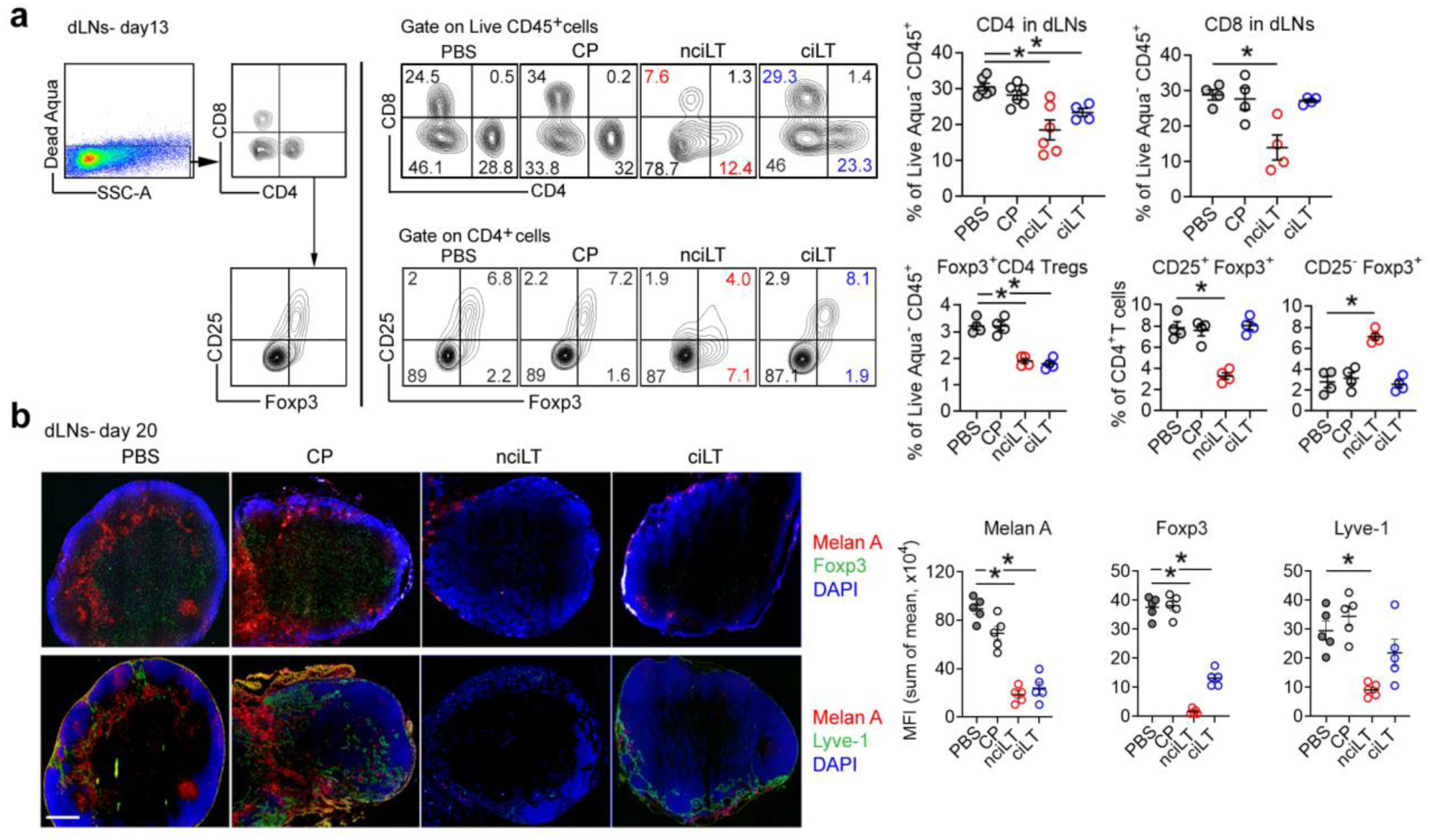
Blockade of LTβR non-classical NFκB signaling reduces CD4 Tregs in dLNs and suppresses melanoma lymphatic metastases. **a**, **b**, Same as in Fig. 7, B16F10 melanoma bearing C57BL/6 mice treated with peritumoral injection of 10 nmol/tumor of nciLT, ciLT, or CP for 5 days. At day 13, dLNs analyzed for CD4, CD8, Foxp3^+^ CD4 Tregs, and T cell IFNγ by flow cytometry (**a**). Draining LNs at day 20 assessed for tumor metastasis (Melan A), Tregs (Foxp3), and lymphangiogenesis (Lyve-1) by immunohistochemistry (**b**). Scale bar: 500 μm. A-B. Mean ± SEM. *p < 0.05 by one-way ANOVA.

## Discussion

In the TME, a high frequency of Tregs is associated with a poor prognosis in many cancers ^37,38^. However, effective approaches of selectively targeting intratumoral Tregs are limited. Beyond the known function of immunosuppression, activated Tregs exert other multifaceted functions such as direct dialogue with endothelial cells to regulate TEM of other immune cells^12,39^. Our current and previous data^11,12^ and published single cell RNA sequencing data^40,41^ have shown that human and mouse CD4 Tregs express the highest level of LTα1β2, while LTβR is predominantly expressed on most tumor cells and endothelial cells^42,43^. Here we demonstrate that contact-dependent tumor LTβR-NFκB-NIK signaling triggered by Treg LTα1β2 plays a critical role in tumor growth and metastases. The TCGA datasets showed high LTβR expressing cancer patients have worse prognosis, suggesting anti-LTβR as a potential therapy. However, both blockade and activation of LTβR have shown anti-tumor efficacy^1,6^. Activation of LTβR with agonist mAb has been shown to induce both tumor apoptosis and promotion^1,6,44^. The seemingly conflicting observations might be due to the bifurcation of LTβR-NFκB signaling. We confirmed that LTβRs on mouse and human melanoma and breast cancer cells signal through classical NFκB-IKKβ-p65 and nonclassical NFκB-NIK-p52. Dissimilar to primary LEC LTβR, which constitutively binds TRAF3 and recruits TRAF2 upon activation, B16F10 LTβR constitutively binds TRAF2 and recruits TRAF3 for activation. nciLT blocked both TRAF2 and TRAF3 binding to the B16F10 LTβR complex, while blocking only TRAF3 binding to LEC LTβR ^3^. Thus, nciLT not only inhibited melanoma LTβR-nonclassical NFκB-NIK-p52 but also inhibited B16F10 LTβR-classical NFκB-IKKβ-p65 signaling.

LTβR-nonclassical NFκB-NIK signaling has been implicated to upregulate metastatic genes and promote cancer cell lymphatic migration in head and neck cancer which has high levels of NIK and RelB expression^45^, however, selective targeting of the LTβR-nonclassical NFκB-NIK pathway has not been studied. Here we employed specific LTβR receptor decoy permeable peptides to examine tumor TEM. LTβR nonclassical NFκB blocking peptide nciLT but not classical blocking peptide ciLT inhibited TEM of most tumors including human melanoma and mouse and human breast cancer. B16F10, 4T1, and A549 TEM or migration responses were also inhibited by ciLT but with lesser efficacy, and these cell type-dependent effects may relate to varied TRAF2 and TRAF3 associations with the LTβR receptor in these cells. Conversely, activation of LTβR promoted tumor TEM. Notably, the presence of LECs significantly promoted tumor cell migration, suggesting LEC derived adhesion or junctional molecules and chemokines are critical. We also observed that blocking LTβR-NIK signaling inhibited tumor cell growth and promoted apoptosis. Classical NFκB signaling is constitutively activated in tumor cells, yet ciLT did not affect tumor growth and produced modest inhibition of B16F10 TEM. LTβR-depleted melanoma cells, in which the entire receptor signaling is ablated, had slower *in vivo* growth and lymphatic metastasis although there was no defect in growth *in vitro*, implicating the importance of environmental cues or intercellular communication in the TME. RNA-seq analysis of LTβR-depleted B16F10 revealed decreased gap junction protein GJA1, PDPN, NOTCH3, and BCAM1 which all regulate cell adhesion and may contribute to the reduced migration and metastasis. Thus, the complexities of LTβR signaling likely account for the diverse effects of receptor blockade or activation. These observations suggest that a more specific approach, as we show here on LTβR signaling pathways, may be more efficacious for tumor therapy.

We found Tregs use LTα1β2 to preferentially stimulate tumor LTβR nonclassical NFκB-NIK signaling to enhance tumor growth and migration. In contrast, neither LTα knockout Tregs nor Teffs enhanced tumor growth or migration. To clarify the role of other related receptor-ligand interactions between Tregs and tumor cells, we analyzed the other ligand of LTβR, LIGHT, however it was not detected on Tregs, while Teffs and B16F10 had negligible LIGHT expression (Extended Data Fig. 7a). Tregs have the highest level of HVEM, the receptor of LIGHT, while B16F10 has minimal HVEM expression, indicating Tregs and Teffs are unlikely to signal B16F10 through LIGHT/HVEM. Tregs but not LTα-deficient Tregs or Teffs induced melanoma LTβR nonclassical NFκB signaling, which was selectively inhibited by nciLT, further evidence that other receptors and ligands of the LTβR family are not involved. Other immune cells, such as B cells and CD8 T cells, also express LTα1β2, but the expression levels are less than that of Tregs in either homeostatic conditions or TME. Similar to Teffs, B cells did not induce LTβR-nonclassical NFkB signaling and had no impact on tumor cell migration. Hence, Tregs are the major source of LTα1β2 to engage LTβR on melanoma or breast cancer cells to promote tumor growth and TEM.

Adoptively transferred WT Tregs but not LTα-deficient Tregs promote tumor growth, and adoptively transferred primary LECs also restore tumor malignancy in LTβR-deficient mice, indicating the critical roles of Treg LTα1β2−LTβR interaction on tumor cells and LECs. Since LTα-deficient Tregs had no defect in immune suppression and viability^11^, failure to suppress immunity was not the cause of their inability to promote tumor growth. Mechanistically, Treg driven LTβR nonclassical NFκB signaling stimulates both tumor and LEC responses that sustain tumor survival, growth, and metastases. In the tumor, this signaling drives expression of angiogenic and immunosuppressive myeloid chemokines CXCL1, CXCL10, and CCL5; protumor interferon-stimulated genes *ISG15* and *IFIT3*; and oncoprotein *USP18*, all of which promote tumor growth and metastases, and are selectively blocked by nciLT but not by ciLT. In LEC, this signaling drives endothelial specific pro-lymphangiogenic SOX18, tumorigenic FLRT2, proangiogenic tumor endothelial molecules Clec14 and Apelin, and myeloid tropic chemokines to promote tumor growth and TEM. The Treg-activated LEC also directly favored tumor growth via cell-cell contact, and nciLT but not ciLT abolished this effect. Additionally, Treg LTα1β2 directly reduced LV and LEC junctional VE-cadherin expression to transform them from continuous “zipper” to discontinuous “button” junctions, promoting CD4 and CD8 egress from the tumor thus impairing intratumoral immunity.

Leveraging in vivo Treg interactions with tumor and LECs, nciLT dramatically suppressed melanoma growth and metastasis with increased intratumoral IFNγ^+^CD8 T cells and reduced tumor CD25^+^CD4^+^ Tregs and MDSC infiltration and angiogenesis. Notably, nciLT reduced SOX18 and FLRT2 on tumor LECs, in line with SOX18-deficient mice having suppressed B16F10 lymphangiogenesis and metastases^46^ and pharmaceutical inhibition of SOX18 reduced breast cancer vascularization and metastases^47^. FLRT2 on tumor endothelial cells forms homophilic interendothelial adhesions to safeguard against oxidative stress, and endothelium-specific deletion of FLRT2 suppressed tumor metastases^35^. Tumor and LEC derived CXCL1 and CXCL10, recruit CXCR2^+^MDSCs^48^ and CXCR3^+^Tregs^49^ (Extended Data Fig.7b), respectively. Thus, suppression of these chemokines by nciLT are likely key attributes for the reduced immune suppressive cells in tumor and enhanced anti-tumor immunity. Since CXCL10 recruit both Tregs and CD8 T cells^50^, blockade of the initial prompt recruitment of Tregs to the tumor bed may play a critical role for tumor regression, which unleashes CD8 proliferation in the TME. In addition to their chemotactic properties, CXCL1 is implicated in tumor growth, angiogenesis, and tumorigenesis^52^. ciLT also enhanced effector T cells and reduced MDSCs in TME, which may contribute to the efficient tumor regression at the early stage. The absence of inhibitory effects of ciLT on the CXCLs chemokines may contribute to later tumor relapse. ciLT also suppressed tumor angiogenesis with less efficiency than nciLT. It is noteworthy that LTβR-classical NFκB-driven *RPL21*, *RPL30*, *MIEN1*, and *NR4a1*, are selectively inhibited by ciLT. These rapidly induced early response protumor genes are overexpressed in various human cancers and proposed as targets for tumor therapy^51–53^. Therefore, ciLT and nciLT combined therapy at selected times may optimize therapeutic efficacy. Together, our data revealed that Treg LTαβ promotes LTβR signaling in tumor cells and LECs. Selectively targeting Tregs or Treg-induced classical and nonclassical LTβR-NFκB signaling on tumor cells and LECs may provide a rational strategy to efficiently prevent tumor growth and metastasis.

## Methods

### Mice

Female C57BL/6J (WT, LTβR^−/−^, LTα^−/−^) (7–10 weeks), were purchased from The Jackson Laboratory (Bar Harbor, ME). Foxp3GFP mice on a C57BL/6 background were from Dr. A. Rudensky (Memorial Sloan Kettering Cancer Center) (Fontenot et al., 2005). LTα^−/−^ mice were crossed with Foxp3GFP mice to generate LTα^−/−^Foxp3GFP mice. All animal experiments were performed in accordance with Institutional Animal Care and Use Committee approved protocols.

### Cell Lines

The mouse melanoma B16F10-eGFP and mouse breast cancer 4T1-eGFP from Imanis Life Sciences (Rochester, MN), B16F10 and Human melanoma A375 from the American Type Culture Collection. were cultured in DMEM with GlutaMAX (4.5 g/L glucose, Invitrogen) containing 10% FCS (Gemini, CA) and 1% Penicillin/Streptomycin (Invitrogen). C57BL/6 mouse (C57-6064L) or human (H-6064L) primary dermal LECs were from Cell Biologics, Inc. (Chicago, IL), and were cultured according to the manufacturer’s instructions in manufacturer-provided mouse endothelial cell medium supplemented with 5% FBS, 2 mM L-glutamine, 100 IU /mL penicillin, vascular endothelial growth factor, endothelial cell growth supplement, heparin, epidermal growth factor, and hydrocortisone.

### Peptides

LTβR decoy peptides were synthesized by GenScript. All peptides were >95% purity, dissolved in endotoxin-free ultrapure water. Peptide reconstitution and usage were described previously^3^. Peptide sequences can be found in Supplementary Table 3.

### Mouse T Cell and B Cell Purification and Culture

Mouse naïve CD4 T and B cells were isolated using mouse CD4 and B cell negative selection kits (Stemcell Technologies, Cambridge, MA). CD4^+^Foxp3GFP^+^ Tregs or CD4^+^Foxp3 GFP^−^ naïve CD4 T cells were sorted from total CD4 T cells from LNs and spleens of WT or LTα^−/−^ Foxp3-GFP mice using a FACS Aria II (BD Biosciences, San Jose, CA) with > 98% purity. The sorted Tregs were cultured overnight at 37 °C in 5% CO2, with medium containing IL-2 (20 ng/ mL, eBioscience, San Diego, CA), plate-bound anti-CD3ɛ mAb (1 μg/ mL, clone 145-2C11, eBioscience), and anti-CD28 mAb (1 μg/mL, clone 37.52, eBioscience), and human TGFβ1 (10 ng/ mL, eBioscience). For effector T cells, the sorted naïve CD4 T cells were cultured with the medium but without human TGFβ1 for 3 days. All cells were cultured in complete RPMI 1640 supplemented with 10% FBS, 1 mM sodium pyruvate, 2 mM L-glutamine, 100 IU/mL penicillin, 100 μg /mL streptomycin, non-essential amino acids and 2 × 10^−5^ M 2-ME (Sigma-Aldrich).

### Human T Cell Purification and Culture

Human Tregs were enriched from peripheral blood mononuclear cells with anti-CD25 microbeads (Miltenyi Biotec, Bergisch Gladbach, Germany), and were sorted via FACSAria as nTreg (CD4^+^CD25^high^CD127^−^CD45RA^+^) and naïve CD4 T cells (CD4^+^CD25^−^CD127^+^CD45RA^+^). The cells were expanded as previously described (Hippen et al., 2011). Briefly, purified nTreg were stimulated with irradiated KT64/86 cells cultured in XVivo-15 (BioWhittaker, Walkersville, MD) media containing 10% human AB serum (Valley Biomedical, Winchester, VA), Pen/Strep (Invitrogen, Carlsbad, CA), N-acetyl cysteine (USP), and recombinant IL-2 (300 IU/mL; Chiron, Emeryville, CA) for 14 days and frozen. When needed, frozen nTreg and naïve CD4 were thawed and re-stimulated with anti-CD3/CD28 mAb-Dynabeads (Life Technologies, Carlsbad, CA) at 1:3 (cell to bead) plus recombinant IL-2 (300 U/mL) for 10 days before assay.

### MTT Viability Assay and Cell Apoptosis Analysis

Cells were plated into 24-well tissue culture plates. After treatment, the cells were washed and followed by 3-hour incubation with 0.5 mg/mL MTT (3-(4, 5-Dimethylthiazol-2-yl)-2,5 diphenyl tetrazolium bromide) (Sigma-Aldrich). Cell apoptosis was measured with Annexin V Apoptosis Detection Kit I (BD Pharmingen) following manufactory instructions.

### Flow Cytometry

Cells were washed and incubated with anti-CD16/32 (clone 93, eBioscience), followed by incubation with fluorescently labeled antibodies (See Supplementary Table 3) for 30 minutes at 4 °C. Cells were then washed and fixed with 4% paraformaldehyde. Data acquisition was performed using an LSR Fortessa flow cytometer (BD Biosciences) and analyzed with FlowJo 8.7 (Tree Star, Inc).

### Immunoblotting

Cells were lysed in buffer containing 20 mM Hepes (pH 7.4), 150 mM NaCl, 10 mM NaF, 2 mM Na_3_VO_4_, 1 mM EDTA, 1 mM EGTA, 0.5% Triton X-100, 0.1 mM DTT, 1 mM PMSF and protease inhibitor cocktail (Roche). Protein in the cell extract was quantified using protein quantification kit (Bio-Rad, Philadelphia, PA) and 10 μg total protein was run on Novex™ WedgeWell™ 4-20% Tris-Glycine Mini Gels (Invitrogen) and transferred to an Immobilon-P membrane (Bio-Rad). Membranes were probed with indicated antibodies (See Supplementary Table 3). Quantification of blots was performed with ImageJ from the National Institutes for Health. Relative intensity of blots was normalized to housekeeping gene GAPDH and presented as fold induction to non-stimulation.

### Quantitative reverse transcription PCR (qRT-PCR) and cDNA microarray analysis

1 μg of total RNA extracted using Trizol reagent (Invitrogen) was reverse transcribed into cDNA with GoScript™ Reverse Transcription System (Promega, Fitchburg, WI). mRNA expression levels were quantified by real-time PCR using SYBR Green Master Mix with an ABI Prism 7900HT (Applied Biosystems, Foster City, CA). Values for specific gene expression were normalized to house-keeping HPRT gene expression and were calculated as: 2^(Ct of HPRT−Ct specific gene). For microarray analysis, samples were further cleaned using an RNeasy Mini Kit (QIAGEN), labeled with Cyanine 3-CTP using a Low Input Quick Amp Labeling Kit (Agilent), and hybridized to an SurePrint G3 Mouse GE Microarray (Agilent). Fluorescence was scanned using a DNA Microarray Scanner (Agilent), and quantified with Feature Extraction ver 10.7.1.1 (Agilent). Expression values for each probe set were calculated using the RMA method with GeneSpring Gx 12.0 software (Agilent). Primers can be found in Supplementary Table 3.

### Bulk RNA sequencing

Total RNA was isolated using the NucleoSpin RNA Plus kit per the manufacturer’s instructions. Messenger RNA was purified and sequenced by Novogene. Reference genome and gene model annotation files were downloaded from Ensembl directly. Index of the reference genome was built using Hisat2 v2.0.5 and paired-end clean reads were aligned to the reference genome using Hisat2 v2.0.5. featureCounts v1.5.0-p3 was used to count the reads numbers mapped to each gene. FPKM (Fragments Per Kilobase of transcript sequence per Millions) of each gene was calculated based on the length of the gene and reads count mapped to this gene. Differential expression analysis of two conditions was performed using the edgeR R package (version 3.22.5). Differential expression analysis of two conditions/groups (two biological replicates per condition) was performed using the DESeq2 R package (1.20.0). The resulting P-values were adjusted using the Benjamini and Hochberg’s approach for controlling the false discovery rate. Genes with an adjusted P-value <=0.05 found by DESeq2 were assigned as differentially expressed. Gene Ontology (GO) enrichment analysis of differentially expressed genes was implemented by the clusterProfiler R package.

### Time-lapse microscopy

WT or LTβR^−/−^B16F10-GFP (5 × 10^4^ cells per transwell) migrating across endothelial monolayers to CCL19 (50 ng/ mL) were visualized by EVOS FL Auto Cell Imaging System (Thermo Fisher Scientific) with a ×20 objective. One image was captured every 5 minutes for 3 hours. Cell tracks were analyzed with Volocity version 6.3 software (Perkin Elmer).

### Tumor Cell Lymphatic Transendothelial Migration

Transmigrations across endothelial cells were described previously (Brinkman et al., 2016; Xiong et al., 2017). Briefly the inverted transwell insert (24-well, Corning International) with 8 μm pore size was coated with 0.2% (w/v) gelatin (Bio-Rad) for 1 hour at 37 °C before loading with 1.0 × 10^5^ primary skin LEC in 100 μL LEC medium. After 2 days, the LEC cell layers were treated with various conditions as noted in the text and figure legends prior to adding 1 × 10^5^ tumor cells in 100 μL to the upper chamber of transwell plate while the lower chamber contained S1P (Avanti Polar Lipids, Inc., Alabaster, AL). All cells or reagents were prepared in IMDM containing transferrin and 0.5% (w/v) fatty acid-free BSA (Gemini, West Sacramento, CA). T cells that migrated to the lower chamber after 3 hours at 37 °C were counted.

### Cell Migration assay

The cells were seeded in a 12-well plate and cultured to 90% confluence. The plates were scratched with a sterile 200 μL pipette tip on the cell monolayer. The cells pretreated with peptides were then cultured for 48 hours. The images were acquired using an ordinary optics microscope (Olympus, Japan) at 0 hour and 48 hours. The migration percentage was calculated according to this formula: (the size of the cell defect at 0 hour − the size of the defect at 48 hours)/the size of the defect at 0 hr.

### Immunohistochemistry

Cell monolayers or tissues were fixed for 20 minutes at 4 °C with 4% (w/v) paraformaldehyde (Affymetrix, Santa Clara, CA), then permeabilized with PBS 0.2% (v/v) Triton X-100 (Sigma-Aldrich) and treated with 4% donkey serum for 30 minutes then incubated with primary antibodies (see Supplementary Table 3.) for overnight at 4°C. The bound antibodies were detected with Alexa Fluor 448, 647 (Cy5), or 546 (Cy3)-conjugated secondary antibodies (Jackson ImmunoResearch, West Grove, PA) for 1 hour at 4 °C. The mounted slides were visualized by fluorescent microscopy (Zeiss LSM 510 Meta and LSM5 Duo). Images were analyzed with Volocity version 6.3 software. Quantification of the junctional VE-cadherin in 60x magnified images of LVs of whole mounted LVs or adherent LECs was performed with ImageJ. Length of zipper junctions and button junctions were measured. Percentage of zipper junction was calculated as: length of zipper junction x 100 / (length of zipper junction + length of button junction) ^54^.

### CRISPR/Cas9 Knockout of LTβR, NIK, and IKKβ in B16F10 and LECs

gRNA designed to target the common exons for all murine LTβR, NIK, and IKKβ isoforms were synthesized as oligo: 5’-AGCCGAGTGCCGCTGTCAGC-3’, 5′-GCTGGCCGCCATCAAGGTTA-3’, and 5′-CGCCATCAAGCAATGCCGAC-3′, respectively and cloned to lentiCRISPRv2 (include Cas9 and puromycin-resistant gene) vector by GenScript. Lentiviruses were produced by co-transfecting HEK293T cells with the packaging plasmids psPAX2 and pMD2.G (Addgene plasmid # 12260 and 12259) and the transfer plasmid LentiCRISPRv2-gRNAs, using Lipofectamine 2000 (Thermo Fisher Scientific). The media was changed to antibiotic-free complete DMEM with 10% FBS after 16 hours. The lentivirus supernatants were collected 24 and 48hours after transfection and filtered through 0.45 mm PES syringe filter (SARSTEDT, Newton, NC). mLEC, B16F10 or B16F10-fluc/eGFP were transduced with the lentiviruses for 3 days, followed by selection medium containing 2 mg/mL puromycin (Sigma-Aldrich) for 3 days. LTβR-lentiCRISPRv2 transduced B16F10-fluc/eGFP cells which have Neomycin and Puromycin resistance, were flow sorted for GFP^+^LTβR^−^ cells. Surviving or sorted cells were expanded and checked with immunoblotting of NIK and IKKβ expression or flow cytometry analysis of LTβR surface expression.

### Tumor treatments

WT or LTβR^−/−^ C57/BL6J mice (7-8 weeks old, female) were subcutaneously injected with 0.5 ×10^6^ WT B16F10 or B16F10-fluc/eGFP tumor cells. 5, 7, 9, and 11 days after tumor inoculation, tumor-bearing mice were peritumoral injected with 20 nmol of CP, nciLT or ciLT. 2 or 9 days after the last peptide treatment, 6 mice from each group were euthanized, and the tumors and dLNs were harvested and analyzed by flow cytometry and immunohistochemistry. Parallel groups of mice were monitored daily for tumor growth. There were 6 to 10 mice in each group. For Treg adoptive transfer assay, induced Tregs differentiated from naïve CD4 T cells from WT or LTα KO Foxp3GFP transgenic mice were FACS sorted before intravenous transfer of 0.5×10^6^ cells per mouse. For LEC adoptive transfer assay, 2×10^5^ primary mouse LECs were intravenously transferred to WT or LTβRKO C57BL/6 mice inoculated with WT B16F10. For pulmonary metastases study, mice were injected via the tail vein with 0.3 ×10^6^ WT B16F10 or B16F10-fluc/eGFP. Lungs were harvested at day 21 after injection.

### Quantification and Statistical Analysis

Numerical data are presented as mean ± SEM. Asterisks mark data statistically different from the controls, with p-values noted in the figure legends. A p-value of <0.05 was considered significant for one-way ANOVA and Student’s *t* tests using Prizm 5 (La Jolla, CA). The number of replicates is noted in the figure legends.

## Data and Code Availability

The RNA-seq and cDNA microarray data are available at the Gene Expression Omnibus. This study did not generate any unique code. Further information and requests for reagents may be directed to, and will be fulfilled by the lead contact Jonathan S. Bromberg

## Acknowledgements

This work was supported by NIH grant R37 AI062765 to JSB, Emerald grant 2022 to JSB and WP, R01 AI126596 and R01 HL141815 to RA, R37 AI34495, R01 HL11879, R01 HL15514, and P01 CA 065493 to BRB and KLH. Dr. Xiaoxuan Fan from UMB-Flow Core Facility for his excellent help for flow cell sorting and data analysis.

## Author Contributions

WP and JSB designed the research. WP, WL, YX, GCZ, SMP, RSO, JS, KH, CMP, DK, MS, YL, AK, LL, AA, VS, and MWS performed the experiments. ACS and SY performed bioinformatic data analysis of the TCGA datasets. WP, JSB, BRB, KLH, RA, PI, JHB, CMJ, and BRB analyzed and discussed the results. WP and JSB wrote the manuscript.

## References

1 Lukashev, M. et al. Targeting the lymphotoxin-beta receptor with agonist antibodies as a potential cancer therapy. Cancer Res 66, 9617–9624, doi:10.1158/0008-5472.CAN-06-0217 (2006).

2 Gommerman, J. L. & Browning, J. L. Lymphotoxin/light, lymphoid microenvironments and autoimmune disease. Nat Rev Immunol 3, 642–655, doi:10.1038/nri1151 (2003).

3 Piao, W. et al. Regulation of T cell afferent lymphatic migration by targeting LTbetaR-mediated non-classical NFkappaB signaling. Nat Commun 9, 3020, doi:10.1038/s41467-018-05412-0 (2018).

4 Wolf, M. J., Seleznik, G. M., Zeller, N. & Heikenwalder, M. The unexpected role of lymphotoxin beta receptor signaling in carcinogenesis: from lymphoid tissue formation to liver and prostate cancer development. Oncogene 29, 5006–5018, doi:10.1038/onc.2010.260 (2010).

5 Keats, J. J. et al. Promiscuous mutations activate the noncanonical NF-kappaB pathway in multiple myeloma. Cancer Cell 12, 131–144, doi:10.1016/j.ccr.2007.07.003 (2007).

6 Hehlgans, T. et al. Lymphotoxin-beta receptor immune interaction promotes tumor growth by inducing angiogenesis. Cancer Res 62, 4034–4040 (2002).

7 Tang, H. et al. Facilitating T Cell Infiltration in Tumor Microenvironment Overcomes Resistance to PD-L1 Blockade. Cancer Cell 30, 500, doi:10.1016/j.ccell.2016.08.011 (2016).

8 Hu, X. et al. Lymphotoxin beta receptor mediates caspase-dependent tumor cell apoptosis in vitro and tumor suppression in vivo despite induction of NF-kappaB activation. Carcinogenesis 34, 1105–1114, doi:10.1093/carcin/bgt014 (2013).

9 Kos, K. et al. Tumor-educated T(regs) drive organ-specific metastasis in breast cancer by impairing NK cells in the lymph node niche. Cell Rep 38, 110447, doi:10.1016/j.celrep.2022.110447 (2022).

10 Kos, K. & de Visser, K. E. The Multifaceted Role of Regulatory T Cells in Breast Cancer. Annu Rev Cancer Biol 5, 291–310, doi:10.1146/annurev-cancerbio-042920-104912 (2021).

11 Brinkman, C. C. et al. Treg engage lymphotoxin beta receptor for afferent lymphatic transendothelial migration. Nat Commun 7, 12021, doi:10.1038/ncomms12021 (2016).

12 Piao, W. et al. Regulatory T Cells Condition Lymphatic Endothelia for Enhanced Transendothelial Migration. Cell Rep 30, 1052–1062 e1055, doi:10.1016/j.celrep.2019.12.083 (2020).

13 Onder, L. et al. Endothelial cell-specific lymphotoxin-beta receptor signaling is critical for lymph node and high endothelial venule formation. J Exp Med 210, 465–473, doi:10.1084/jem.20121462 (2013).

14 Zhai, Y. et al. LIGHT, a novel ligand for lymphotoxin beta receptor and TR2/HVEM induces apoptosis and suppresses in vivo tumor formation via gene transfer. J Clin Invest 102, 1142–1151, doi:10.1172/JCI3492 (1998).

15 Leong, S. P., Naxerova, K., Keller, L., Pantel, K. & Witte, M. Molecular mechanisms of cancer metastasis via the lymphatic versus the blood vessels. Clin Exp Metastasis 39, 159–179, doi:10.1007/s10585-021-10120-z (2022).

16 Kohli, K., Pillarisetty, V. G. & Kim, T. S. Key chemokines direct migration of immune cells in solid tumors. Cancer Gene Ther 29, 10–21, doi:10.1038/s41417-021-00303-x (2022).

17 Yang, H. et al. Sox17 promotes tumor angiogenesis and destabilizes tumor vessels in mice. J Clin Invest 123, 418–431, doi:10.1172/JCI64547 (2013).

18 Grimm, D. et al. The role of SOX family members in solid tumours and metastasis. Semin Cancer Biol 67, 122–153, doi:10.1016/j.semcancer.2019.03.004 (2020).

19 Chen, H., Xu, C., Jin, Q. & Liu, Z. S100 protein family in human cancer. Am J Cancer Res 4, 89–115 (2014).

20 Acharyya, S. et al. A CXCL1 paracrine network links cancer chemoresistance and metastasis. Cell 150, 165–178, doi:10.1016/j.cell.2012.04.042 (2012).

21 Cartier, A. & Hla, T. Sphingosine 1-phosphate: Lipid signaling in pathology and therapy. Science 366, doi:10.1126/science.aar5551 (2019).

22 Xiong, Y. et al. A robust in vitro model for trans-lymphatic endothelial migration. Sci Rep 7, 1633, doi:10.1038/s41598-017-01575-w (2017).

23 Wang, N. et al. CXCL1 derived from tumor-associated macrophages promotes breast cancer metastasis via activating NF-kappaB/SOX4 signaling. Cell Death Dis 9, 880, doi:10.1038/s41419-018-0876-3 (2018).

24 Kim, M., Choi, H. Y., Woo, J. W., Chung, Y. R. & Park, S. Y. Role of CXCL10 in the progression of in situ to invasive carcinoma of the breast. Sci Rep 11, 18007, doi:10.1038/s41598-021-97390-5 (2021).

25 Aldinucci, D. & Colombatti, A. The inflammatory chemokine CCL5 and cancer progression. Mediators Inflamm 2014, 292376, doi:10.1155/2014/292376 (2014).

26 Bolado-Carrancio, A. et al. ISGylation drives basal breast tumour progression by promoting EGFR recycling and Akt signalling. Oncogene 40, 6235–6247, doi:10.1038/s41388-021-02017-8 (2021).

27 Pidugu, V. K., Pidugu, H. B., Wu, M. M., Liu, C. J. & Lee, T. C. Emerging Functions of Human IFIT Proteins in Cancer. Front Mol Biosci 6, 148, doi:10.3389/fmolb.2019.00148 (2019).

28 Hong, B. et al. USP18 is crucial for IFN-gamma-mediated inhibition of B16 melanoma tumorigenesis and antitumor immunity. Mol Cancer 13, 132, doi:10.1186/1476-4598-13-132 (2014).

29 Kang, J. et al. Ribosomal proteins and human diseases: molecular mechanisms and targeted therapy. Signal Transduct Target Ther 6, 323, doi:10.1038/s41392-021-00728-8 (2021).

30 Kpetemey, M. et al. MIEN1, a novel interactor of Annexin A2, promotes tumor cell migration by enhancing AnxA2 cell surface expression. Mol Cancer 14, 156, doi:10.1186/s12943-015-0428-8 (2015).

31 Hibino, S. et al. Inhibition of Nr4a Receptors Enhances Antitumor Immunity by Breaking Treg-Mediated Immune Tolerance. Cancer Res 78, 3027–3040, doi:10.1158/0008-5472.CAN-17-3102 (2018).

32 Sidibe, A. & Imhof, B. A. VE-cadherin phosphorylation decides: vascular permeability or diapedesis. Nat Immunol 15, 215–217, doi:10.1038/ni.2825 (2014).

33 Futterer, A., Mink, K., Luz, A., Kosco-Vilbois, M. H. & Pfeffer, K. The lymphotoxin beta receptor controls organogenesis and affinity maturation in peripheral lymphoid tissues. Immunity 9, 59–70, doi:10.1016/s1074-7613(00)80588-9 (1998).

34 Young, N. et al. Effect of disrupted SOX18 transcription factor function on tumor growth, vascularization, and endothelial development. J Natl Cancer Inst 98, 1060–1067, doi:10.1093/jnci/djj299 (2006).

35 Ando, T. et al. Tumor-specific interendothelial adhesion mediated by FLRT2 facilitates cancer aggressiveness. J Clin Invest 132, doi:10.1172/JCI153626 (2022).

36 Olbromski, M., Podhorska-Okolow, M. & Dziegiel, P. Role of the SOX18 protein in neoplastic processes. Oncol Lett 16, 1383–1389, doi:10.3892/ol.2018.8819 (2018).

37 Petersen, R. P. et al. Tumor infiltrating Foxp3+ regulatory T-cells are associated with recurrence in pathologic stage I NSCLC patients. Cancer 107, 2866–2872, doi:10.1002/cncr.22282 (2006).

38 Saleh, R. & Elkord, E. Acquired resistance to cancer immunotherapy: Role of tumor-mediated immunosuppression. Semin Cancer Biol 65, 13–27, doi:10.1016/j.semcancer.2019.07.017 (2020).

39 Ring, S. et al. Regulatory T Cells Prevent Neutrophilic Infiltration of Skin during Contact Hypersensitivity Reactions by Strengthening the Endothelial Barrier. J Invest Dermatol 141, 2006–2017, doi:10.1016/j.jid.2021.01.027 (2021).

40 He, M. X. et al. Transcriptional mediators of treatment resistance in lethal prostate cancer. Nat Med 27, 426–433, doi:10.1038/s41591-021-01244-6 (2021).

41 Wu, S. Z. et al. A single-cell and spatially resolved atlas of human breast cancers. Nat Genet 53, 1334–1347, doi:10.1038/s41588-021-00911-1 (2021).

42 Wu, S. Z. et al. Cryopreservation of human cancers conserves tumour heterogeneity for single-cell multi-omics analysis. Genome Med 13, 81, doi:10.1186/s13073-021-00885-z (2021).

43 Zilionis, R. et al. Single-Cell Transcriptomics of Human and Mouse Lung Cancers Reveals Conserved Myeloid Populations across Individuals and Species. Immunity 50, 1317–1334 e1310, doi:10.1016/j.immuni.2019.03.009 (2019).

44 Daller, B. et al. Lymphotoxin-beta receptor activation by lymphotoxin-alpha(1)beta(2) and LIGHT promotes tumor growth in an NFkappaB-dependent manner. Int J Cancer 128, 1363–1370, doi:10.1002/ijc.25456 (2011).

45 Das, R. et al. Lymphotoxin-beta receptor-NIK signaling induces alternative RELB/NF-kappaB2 activation to promote metastatic gene expression and cell migration in head and neck cancer. Mol Carcinog 58, 411–425, doi:10.1002/mc.22938 (2019).

46 Duong, T. et al. Genetic ablation of SOX18 function suppresses tumor lymphangiogenesis and metastasis of melanoma in mice. Cancer Res 72, 3105–3114, doi:10.1158/0008-5472.CAN-11-4026 (2012).

47 Overman, J. et al. Pharmacological targeting of the transcription factor SOX18 delays breast cancer in mice. Elife 6, doi:10.7554/eLife.21221 (2017).

48 Highfill, S. L. et al. Disruption of CXCR2-mediated MDSC tumor trafficking enhances anti-PD1 efficacy. Sci Transl Med 6, 237ra267, doi:10.1126/scitranslmed.3007974 (2014).

49 Moreno Ayala, M. A., et al. CXCR3 expression in regulatory T cells drives interactions with type I dendritic cells in tumors to restrict CD8(+) T cell antitumor immunity. Immunity 56, 1613–1630 e1615, doi:10.1016/j.immuni.2023.06.003 (2023).

50 Ozga, A. J., Chow, M. T. & Luster, A. D. Chemokines and the immune response to cancer. Immunity 54, 859–874, doi:10.1016/j.immuni.2021.01.012 (2021).

51 Guo, H. et al. NR4A1 regulates expression of immediate early genes, suppressing replication stress in cancer. Mol Cell 81, 4041–4058 e4015, doi:10.1016/j.molcel.2021.09.016 (2021).

52 Kushwaha, P. P., Gupta, S., Singh, A. K. & Kumar, S. Emerging Role of Migration and Invasion Enhancer 1 (MIEN1) in Cancer Progression and Metastasis. Front Oncol 9, 868, doi:10.3389/fonc.2019.00868 (2019).

53 Li, C. et al. RPL21 siRNA Blocks Proliferation in Pancreatic Cancer Cells by Inhibiting DNA Replication and Inducing G1 Arrest and Apoptosis. Front Oncol 10, 1730, doi:10.3389/fonc.2020.01730 (2020).

54 Zhang, F. et al. Lacteal junction zippering protects against diet-induced obesity. Science 361, 599–603, doi:10.1126/science.aap9331 (2018).

