## supplemental figures for "Regulatory T cells crosstalk with tumor and endothelium through lymphotoxin signaling"

**Supplemental Information**

**Supplementary table 1. LTβR expression in various cancer cells**

| Cancer type | Cell line | Species | LTβR (MFI) |
|---|---|---|---|
| Breast | 66.1 | mouse | 7328 *** |
| | 410 | mouse | 4360 *** |
| | 4T1 | mouse | 2747 *** |
| | MDA-MB-231, clone 304 (Low metastatic) | human | 1285 ** |
| | MDA-MB-231, clone 308 (High metastatic) | human | 1527 *** |
| | MDA-MB-468 | human | 636 * |
| | T47D | human | 538 * |
| Sarcoma | KPI30 | mouse | 2436 *** |
| | HT1080 | human | 7554 *** |
| Lung | LLC3 | mouse | 1324 ** |
| | A549 | human | 1747 *** |
| Skin | B16F10 | mouse | 3312 *** |
| | SCC13 | human | 43 * |
| | A375 | human | 1812 *** |
| Ovarian | ID-8 | human | 5426 *** |
| | CaoV3 | human | 8280 *** |
| | ES-2 | human | 5049 *** |
| B cell lymphoma | M12.4 | mouse | 494 * |

*: MFI<600, low; **: MFI 600-1500, medium; ***: MFI>1500, high

**Supplementary table 2. Inhibition of tumor TEM by LTβR blocking peptides in various cancer cells**

| Cancer type | Cell line | Species | Peptide inhibition of in vitro TEM |
|---|---|---|---|
| Breast cancer | 66.1 | mouse | nciLT |
| | 410 | mouse | nciLT |
| | MDA-MB-231, clone 308 (high metastatic) | human | nciLT |
| | MDA-MB-231, clone 6 (low metastatic) | human | nciLT |
| Sarcoma | KPI30 | mouse | nciLT |
| | HT1080 | human | nciLT |
| | A549 | human | nciLT |
| Melanoma | B16F10 | mouse | nciLT or ciLT |
| | A375 | human | nciLT |
| Lung cancer | LLC1 | mouse | nciLT or hciLT |
| | A549 | human | nciLT or hciLT |

**Supplementary table 3. Source of antibodies, primer sequencies, and peptide sequences.**

| Antibodies Dilution | SOURCE | IDENTIFIER and |
|---|---|---|
| Anti-NFκB2 p100/p52 | Cell Signaling Technology | Cat# 4882; RRID:AB_10828354 |
| Anti-phospho-NFκBp65 (Ser536) | Cell Signaling Technology | Cat# 3033; RRID:AB_331284 |
| *A*nti-NFκBp65 (D14E12) | Cell Signaling Technology | Cat# 8242; RRID:AB_10859369 |
| Anti-GAPDH (14C10) | Cell Signaling Technology | Cat# 2118; RRID:AB_561053 |
| Anti-phospho-I KK α/β(Ser176/180) | Cell Signaling Technology | Cat# 2697; RRID:AB_2079382 |
| Anti-IKKβ (D30C6) | Cell Signaling Technology | Cat# 8943; RRID:AB_11024092 |
| Anti-TRAF2 | Cell Signaling Technology | Cat# 4712; RRID:AB_2209848 |
| Anti-TRAF3 | Cell Signaling Technology | Cat# 4729; RRID:AB_2209417 |
| Anti-mouse LTβR (5G11b) | Biolegend | Cat# 134402; RRID:AB_1659177 |
| Anti-mouse LTβR poly clonal Ab | ThermoFisher | Cat# PA5-75283, |
| Donkey anti-rabbit IgG HL-HPRT | Cell Signaling Technology | Cat# 7074; RRID:AB_2099233 |
| Donkey anti-mouse IgG HL-HPRT | Cell Signaling Technology | Cat# 7076; RRID:AB_330924 |
| Donkey anti-rat IgG HL-HPRT | Cell Signaling Technology | Cat# 7077; RRID:AB_10694715 |
| Goat anti-rat IgG, Light-Chain Specific -HPRT | Cell Signaling Technology | Cat# 98164; RRID:AB_2936818 |
| Anti-mouse LTβR (eBio3C8), Functional Grade | ThermoFisher | Cat# 16-5671-82; RRID:AB_763451 |
| Anti-CD28 (37.51), Functional Grade | ThermoFisher | Cat# 16-0281-82; RRID:AB_468921 |
| Anti-CD3ε (145-2C11) | ThermoFisher | Cat# 14-0031-86; |
| Anti-human LTβR-PE (31G4D8) | Biolegend | Cat# 322008; RRID:AB_2139070 |
| Anti-mouse LTβR-PE (3C8) | ThermoFisher | Cat# 12-5671-80 |
| Anti-mouse LTβR-PE (5G11) | Biolegend | Cat# 134403; RRID:AB_1659180 |
| *A*nti-human CD258 APC | Biolegend | Cat# 318709; RRID:AB_2728269 |
| Anti-mouse LIGHT AF 647 | R&D systems | Cat# FAB17942R; N/A |
| Anti-human HVEM (122) PE | Biolegend | Cat# 318805; RRID:AB_2203704 |
| Anti-mouse HVEM (1b18) APC | Biolegend | Cat# 136305; RRID:AB_2203829 |
| *A*nti-CD45-BV605 | BD Biosciences | Cat# 567459, RRID:AB_2916603 |
| Anti-mouse CD4 (Gk1.5) AF700 | Biolegend | Cat# 100429; RRID:AB-493699 |
| Anti-mouse CD8a (53-6.7) APC/Cy7 | Biolegend | Cat# 100713; RRID:AB_312753 |
| *A*nti-mouse CD25 (A18246a) PE | Biolegend | Cat# 113703; RRID:AB_2927943 |
| *A*nti-CD25-PE (PC61.5) | ThermoFisher | Cat# 12-0251-83 |

| | | |
|---|---|---|
| *A*nti-Foxp3 (MF-14) BV421 | Biolegend | Cat#126419; RRID:AB_2565933 |
| *A*nti-IFNγ-PerCP/Cy5 | Biolegend | Cat#505821 |
| *An*ti-mouse/humanB220 (Ra3-6b2) APC | Biolegend | Cat# 103211; RRID:AB_312997 |
| Anti-mouse/human CD11b (M1/70) Pacific Blue | Biolegend | Cat# 101223; RRID:AB_755985 |
| Anti-mouse F4/80 (Bm8) PerCP/Cy5.5 | Biolegend | Cat#123127; RRID:AB_893496 |
| *A*nti-Ly-6G (1A8) PE | ThermoFisher | Cat# 12-9668-82; RRID:AB_2572720 |
| Anti-Melan-A/MART-1 (A103) | ThermoFisher | Cat# MA1-34864, RRID:AB_1956105 |
| Anti-mouse CD31(PECAM-1 | BD Biosciences | Cat# 561813; |
| Anti-mouse Lyve-1 | R&D systems | Cat# AF2125; RRID:AB_2297188 |
| Anti-mouse Lyve-1 | Fitzgerald | Cat# 70R-LR003; RRID:AB_1287923 |
| Anti-mouse FLRT2 | ThermoFisher | Cat# PA5 109729;RRID:AB_2855140 |
| Anti-human/mouse SOX18 | ThermoFisher | Cat# PA5-40640; RRID:AB_2608888. |
| Anti-mouse Clec14 | ThermoFisher | Cat# PA5-111267 RRID:AB_2856677 |
| Donkey anti-Rabbit IgG AF647 | ThermoFisher | Cat# A-31573; RRID:AB_2536183. 1:400 |
| Donkey anti-Rabbit IgG AF647 | Jackson Immuno Research | Cat# 711-605-152; RRID:AB_2492288. 1:400 |
| Donkey anti-Rabbit IgG Cy™3 | Jackson Immuno Research | Cat# 711-165-152; RRID:AB_2307443. 1:400 |
| Donkey anti-Rat IgG AF647 | Jackson Immuno Research | Cat# 712-605-153; RRID:AB_2340694. 1:400 |
| Donkey anti-Rat IgG Cy™3 | Jackson Immuno Research | Cat# 712-165-153; RRID:AB_2340667. 1:400 |
| Donkey anti-Goat IgG Cy™3 | Jackson Immuno Research | Cat# 705-165-147; RRID:AB_2307351. 1:400 |
| Donkey anti-Goat IgG AF647 | Jackson Immuno Research | Cat# 705-605-003; RRID:AB_2340436. 1:400 |
| anti-Mouse IgG Alexa Flour 647 | Jackson Immuno Research | Cat# 715-605-150. 1:400 |
| Rat IgG1 (MOPC21) | BioXCell | Cat# BE0083. |
| Anti-mouse IgG1-BV421(RMG1) | Biolegend | Cat# 406616; RRID:AB_2562234 |
| *A*nti-mouse CCL5 (2E9) PE | Biolegend | Cat#149103. 1:200 |
| Anti- mouse CXCL10 | R&D systems | Cat#MAB466. 1:200 |
| Anti- mouse CXCL10 Biotin | R&D systems | Cat #: BAF466 |
| Streptavidin AF700 | ThermoFisher | Cat#: S21383 |
| *A*nti-human CXCL1 Alexa Flour 647 | BD Bioscience | Cat#566579. 1:200 |
| Anti-human CXCL10 PerCP/Cy5.5 (J034d6) | Biolegend | Cat#519509. 1:200 |
| **Oligonucleotides** | **Forward (5'-3')** | **Reverse (5'-3')** |
| Primer: mCCL5 | CAAGTGCTCCAATCTTGCAGTC | TTCTCTGGGTTGGCACACAC |
| Primer: mCXCL1 | GGGCGCCTATCGCCAAT | ACCTTCAAGCTCTGGATGTTC TT |

| | | |
|---|---|---|
| Primer: mCXCL5 | CTGGATCCAGAAGCTCCTGT | CGAGTGCATTCCGCTTAGCT |
| Primer: mCXCL10 | GCCGTCATTTTCTGCCTCA | CGTCCTTGCGAGAGGGATC |
| Primer: IFIT3 | CCTACTCCGTGAAGTCTAGGG | AGTGTGAAGGTTTTGAGCGT |
| Primer: USP18 | CCCTCATGGTCTGGTTGGTT | GTTCAGCTCCTTCCACGCTT |
| Primer: ISG15 | TCTGACTGTGAGAGCAAGCAG | ACCTTTAGGTCCCAGGCCATT |
| Primer: RPL21 | CGCAGCCATCTTCCAGTAAC | CGCATGTATGTGGCCAAAGG |
| Primer: MIEN1 | GGAGGAGTATCCGGGCATTG | GGTTCTCCATTGCTGGCTCT |
| Primer: FLRT2 | GGAGACAAGGCTGCCAGATT | ATGCAGTGGTCCTCCATGTG |
| Primer: SOX18 | CGACTGGCGCAACAAAATCC | GTTCAGCTCCTTCCACGCTT |
| Primer: Clec14a | ATGCAGTGGTCCTCCATGTG | GAAGACCACCGTGGAAGAG |
| Primer: APLN | CTGGGTTTAGTGAAGCGGGT | TTAGGGACTATTCCGGGCCA |
| Primer: HPRT | TGAAGAGCTACTGTAATGATCAGTCAAC | AGCAAGCTTGCAACCTTAACCA |
| **Peptides** | **Sequences** | |
| nciLT | RQIKIWFQNRRMKWKKTGNIYIYNGPVLG | |
| ciLT | RQIKIWFQNRRMKWKKTPEEGAPGP | |
| CP | RQIKIWFQNRRMKWKKGEHGQVAHGA | |

**Extended Data Fig. 1: LT and LTβR gene expression in major cell component of human cancers.**

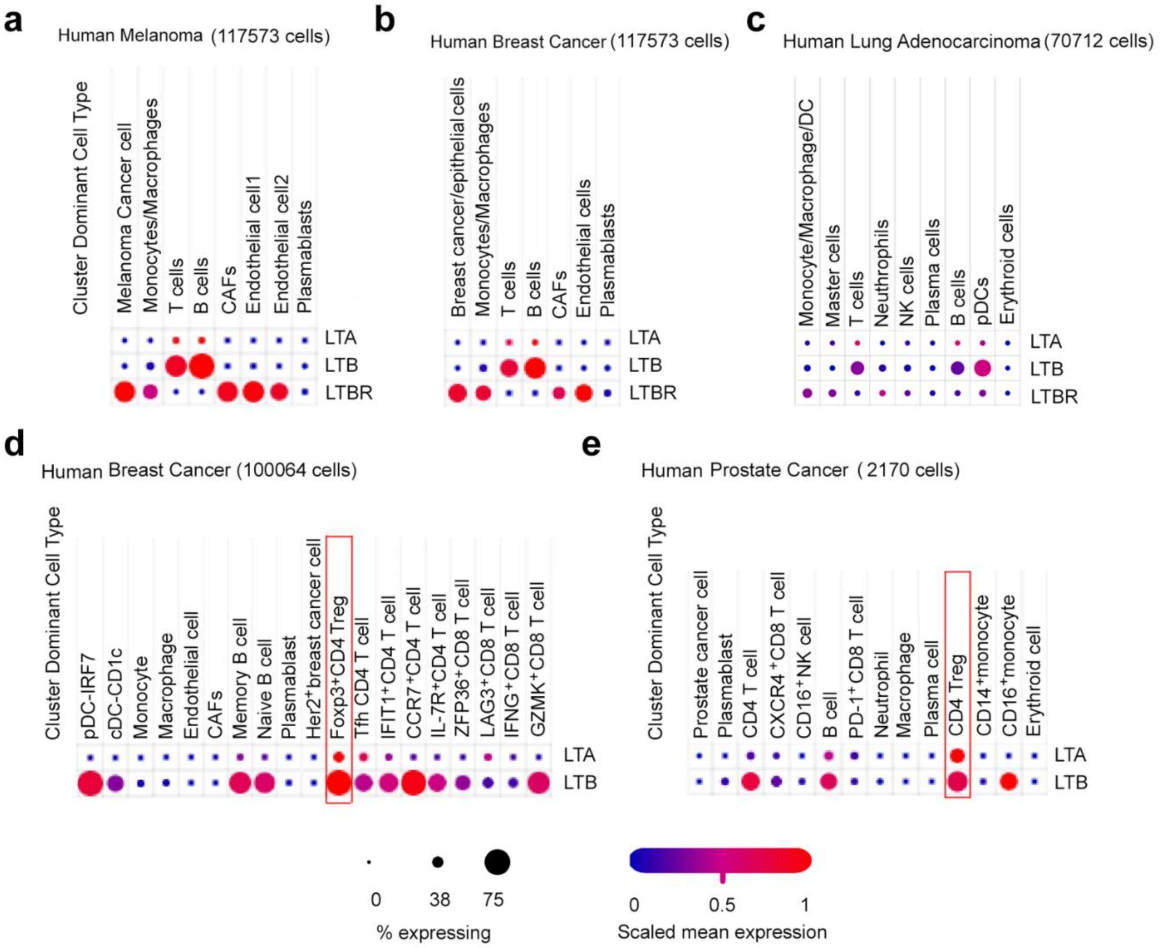


(a-c) Dot plots of LTA, LTB, and LTBR gene expression in major cell components of human melanoma[1] (a), breast cancer[1] (b), and lung adenocarcinoma[2] (c) from published single cell RNA sequencing data. (d-e) LTA and LTB gene expression in breast cancer[3] (d) and prostate cancer[4] (e) immune cell subsets of TILs from published single cell RNA sequencing data.

**Extended Data Fig. 2: Enrichment analysis of DEGs in CRISPR/Cas9 LTβR KO B16F10 cells.**

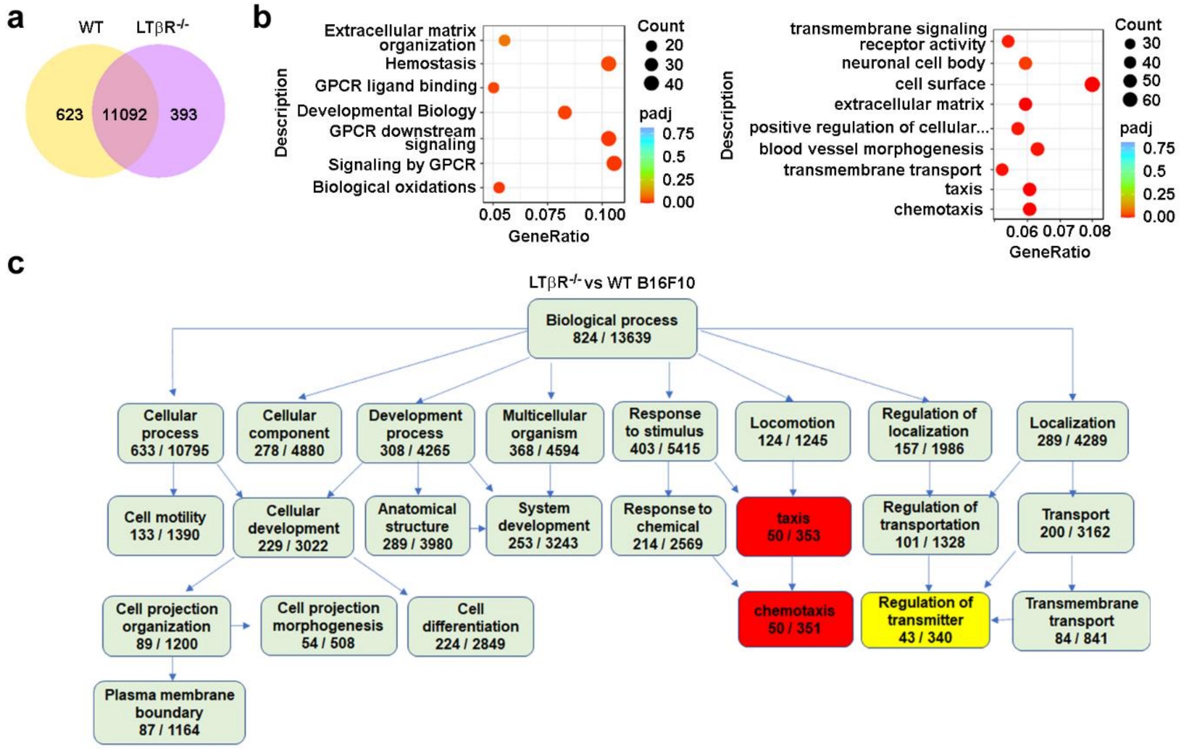


(a) Venn diagrams listing numbers of DEGs in CRISPR/Cas9 LTβR KO (LTβR$^{-/-}$) vs WT B16F10. (b) Gene ontology (GO) analysis of DEGs for molecular and biological functions. GO terms enriched in LTβR$^{-/-}$ vs WT B16F10. GO terms with corrected P value < 0.05 considered significantly enriched by DEGs. (c) GO diagram of LTβR dependent biological processes.

**Extended Data Fig. 3: CXCL1, CXCL10 and CCL5 expression are abolished in NIK-depleted but not IKKβ depleted cells B16F10. Human Treg induced CXCL1 and CXCL10 production in human melanoma and LECs are selectively suppressed by LTβR-nonclassical NFkB blockade.**

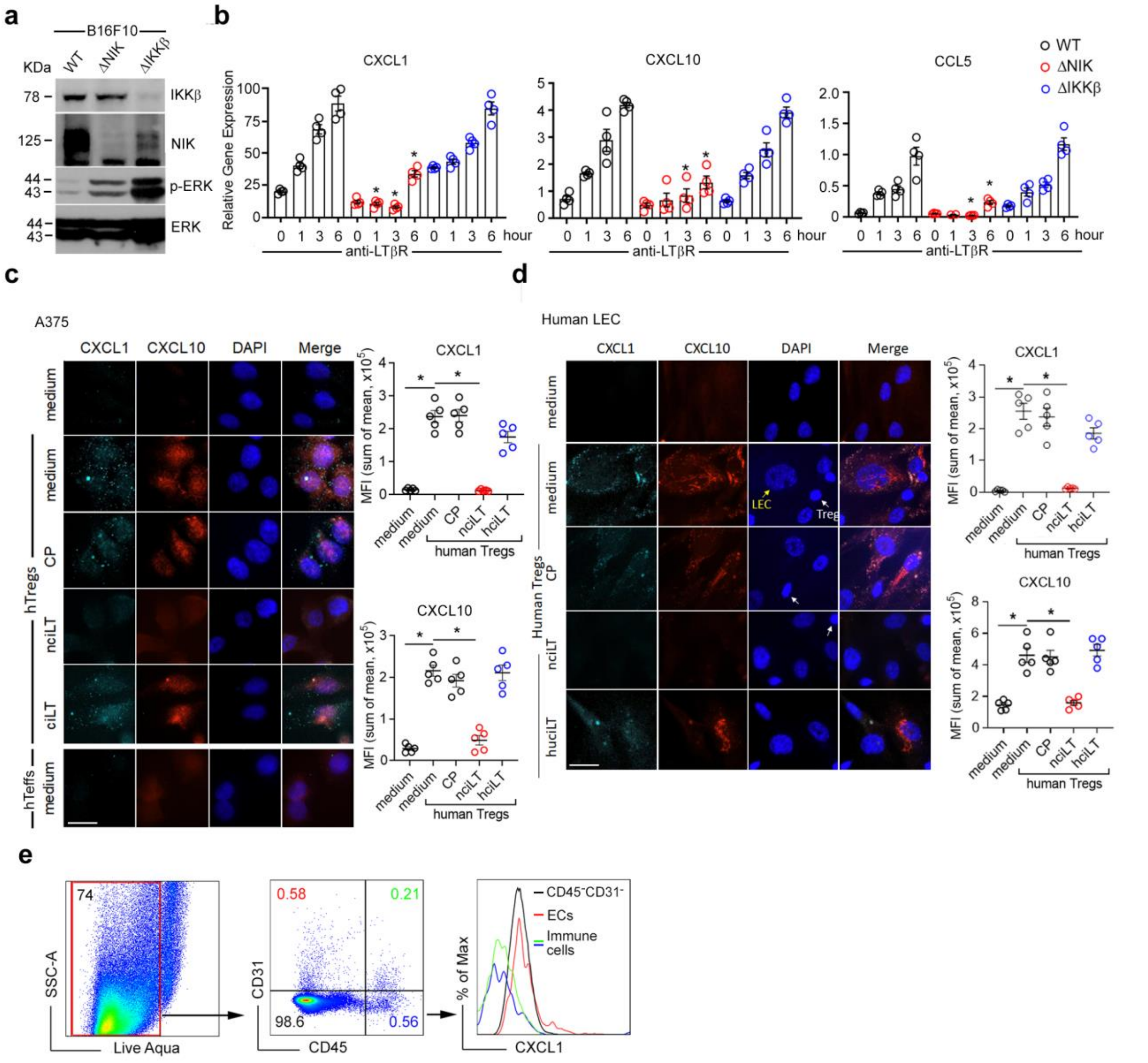

(a) Immunoblot of IKKβ, NIK, phospho-ERK, and total ERK expression in CRISPR/Cas9 IKKβ knock out (ΔIKKβ) and CRISPR/Cas9 NIK knock out (ΔNIK). (b) Real-time PCR of CXCL1, CXCL10, and CCL5 in WT, ΔNIK, and ΔIKKβ B16F10 stimulated with 2 μg/mL agonist anti-LTβR Ab (3C8) for 1, 3, and 6 hours. (c-d) Immunohistochemistry of CXCL1 and CXCL10 expression in human melanoma A375 (c) and human LECs (d). 3x10[4] A375 or human LECs pretreated with 20 μM of indicated peptides for 1 hour, washed and cocultured with $1x10^5$ human Tregs or Teffs for 6 hours. Magnification 60x; scale bar:25 μm. Mean ± SEM. *$p < 0.05$ by one-way ANOVA. (e) $CD45^+$ immune cells, $CD45^-CD31^-$ tumor cells including fibroblast cells, and $CD45^-CD31^+$ endothelial cells (ECs) from transplanted mouse melanoma after 13 days assessed for intracellular CXCL1 expression.

**Extended Data Fig. 4: Tregs exclusively induce LTβR- nonclassical NFκB signaling in tumor cells and promote TEM.**

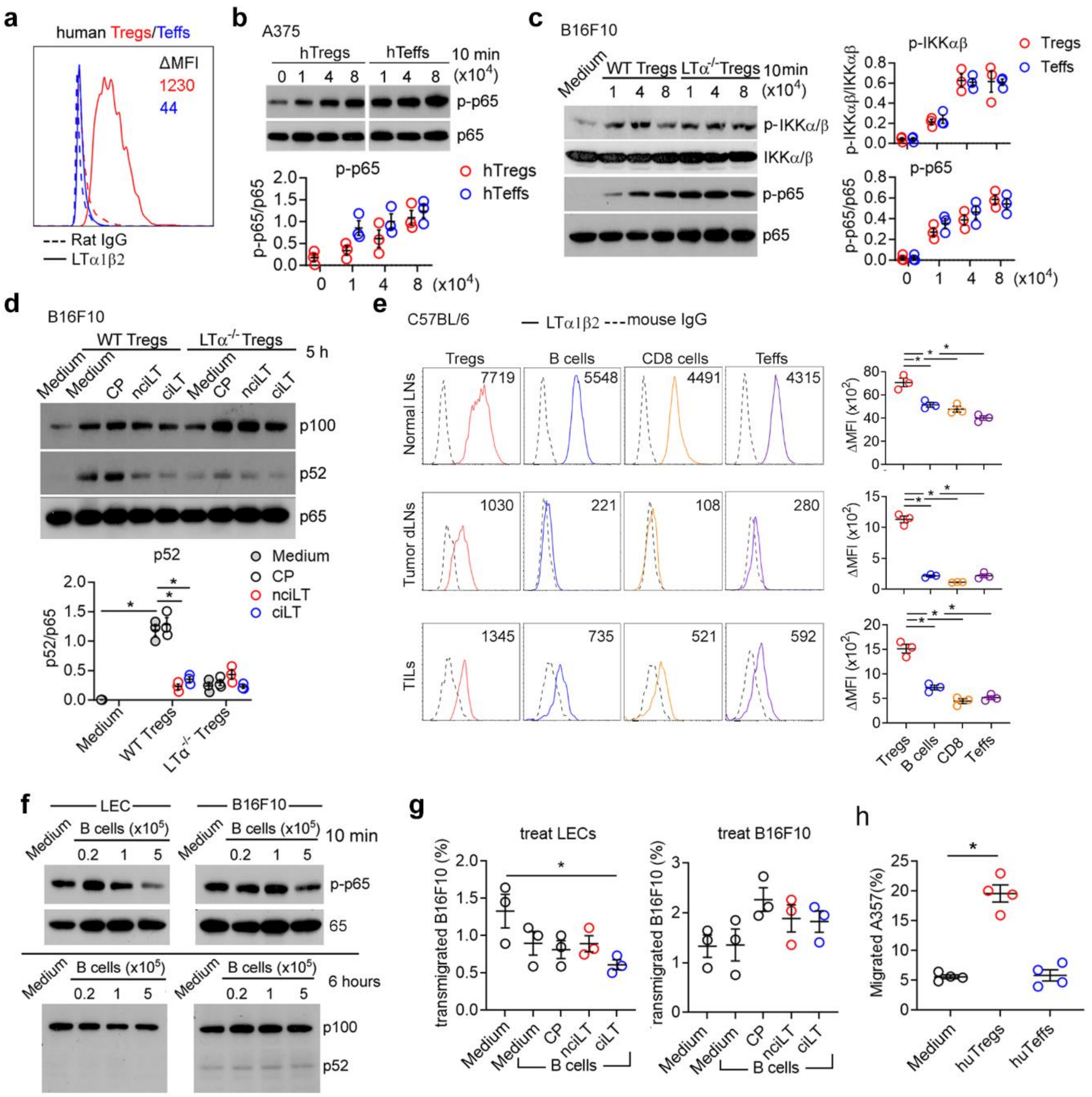

(a) LTα1β2 surface expression on human Tregs or Teffs. ΔMFI shown. (b) Immunoblot of phosphorylated p65 in human melanoma A375 cells co-cultured with various doses of human Tregs or Teffs for 10 minutes. (c) Immunoblot of phosphorylated IKKα/β and p65 in B16F10 co-cultured with various doses of WT or LTα-deficient (LTα$^{-/-}$) Tregs for 10 minutes. (d) Immunoblot of p100/p52 in B16F10 pretreated with 20 μM CP, nciLT, or ciLT for 1 hour prior to co-culture with WT or LTα$^{-/-}$ Tregs for 5 hours. (e) LTα1β2 surface expression on mouse Tregs, B cells, CD8 T cells, and Teffs from normal LNs or B16F10 melanoma dLNs and TILs. ΔMFI shown. (f) Immunoblot of phosphorylated p65 and p100/p52 in B16F10 co-cultured with indicated numbers of B cells for 10 minutes and 6 hours, respectively. (g) B16F10-GFP or LECs separately pretreated with 20 μM of indicated peptides for 30 min, then separately co- cultured for 16 hours with $5x10^5$ B cells, then B16F10 TEM assessed. (h) A375 co-cultured with human Tregs or Teffs for 5 hours prior to 16 hours transwell migration assay. Data representative of 3 experiments. Mean ± SEM. *$p < 0.05$ by one-way ANOVA.

**Extended Data Fig. 5: LEC secretomes induced by Treg co-culture did not affect tumor growth; direct physical contact between B16F10 and WT but not LT$\alpha^{-/-}$ Tregs promotes tumor growth.**

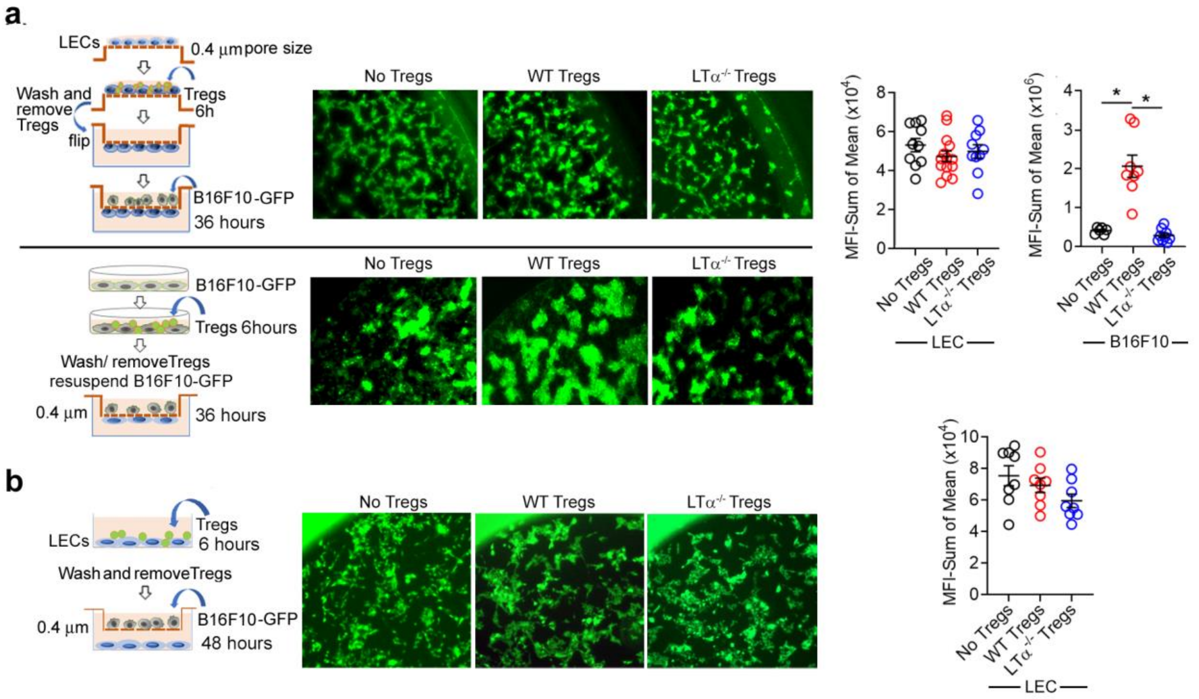


(a-b) Indirect co-culture of B16F10 and LECs. $3x10^4$ LECs coated either on 0.4 μm pore transwell (a) or on 24-well TC plates (b) for 48 hours, then cocultured with WT or LT$\alpha^{-/-}$Tregs for 6 hours, then Tregs removed. $3x10^3$ B16F10s loaded into upper chamber allowing for communication with LECs via cell secretomes, and co-cultured for 36 hours (a, upper panel) or 48 hours (b). B16F10 directly cocultured with WT or LT$\alpha^{-/-}$ Tregs for 6 hours allowing for communication though their surface receptors, then Tregs removed and B16F10 cultured for 36 hours (a, lower panels).

**Extended Data Fig.6: Tumor immune cell infiltration in melanoma-bearing LTβR$^{-/-}$mice**

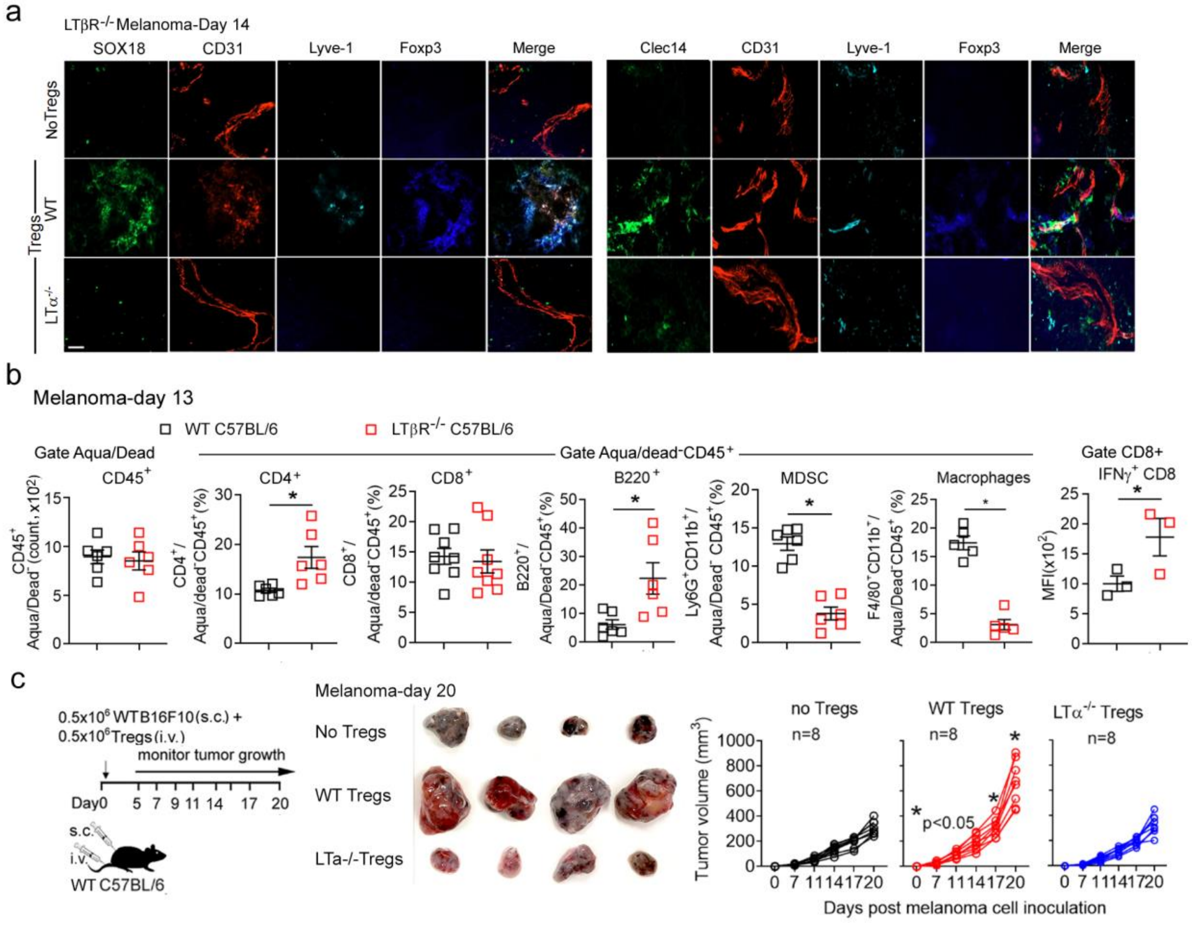


(a) Immunohistochemistry of Sox18, Clec14, and Foxp3 expression on Lyve-1$^+$CD31$^+$LECs within melanoma at day 14 after 5x10$^5$ WT B16F10 inoculation together with 5 x10$^5$ WT or LTα$^{-/-}$Treg intravenous transfer. Magnification 20x; scale bar:42 μm. (b**)** 0.5 x10$^6$ WT B16F10 intradermally injected into WT C57BL6 or LTβR$^{-/-}$ C57BL/6 mice. At day13, tumor CD4 and CD8 T cells, B220$^+$ B cells, Ly6G$^+$CD11b$^+$ MDSCs, F4/80$^+$CD11b$^+$ macrophages, IFNγ+ CD8, analyzed by flow cytometry. (c**)** WT C57BL/6 mice injected intradermally with 0.5 x10$^6$ WT B16F10 and intravenously with or without 1 x10$^6$ WT Tregs or LTα$^{-/-}$ Tregs. Experimental scheme, tumors at day 20, and melanoma growth shown. Data represent 2 independent experiments. with 8 mice in each group. Mean ± SEM. *p < 0.05 by one-way ANOVA.

**Extended Data Fig. 7: Expression of LIGHT, HVEM, and LTβR on B16F10, Tregs, and Teffs. CXCR3⁺Tregs and CXCR2⁺MDSCs in B16F10 melanoma TIL and dLNs**

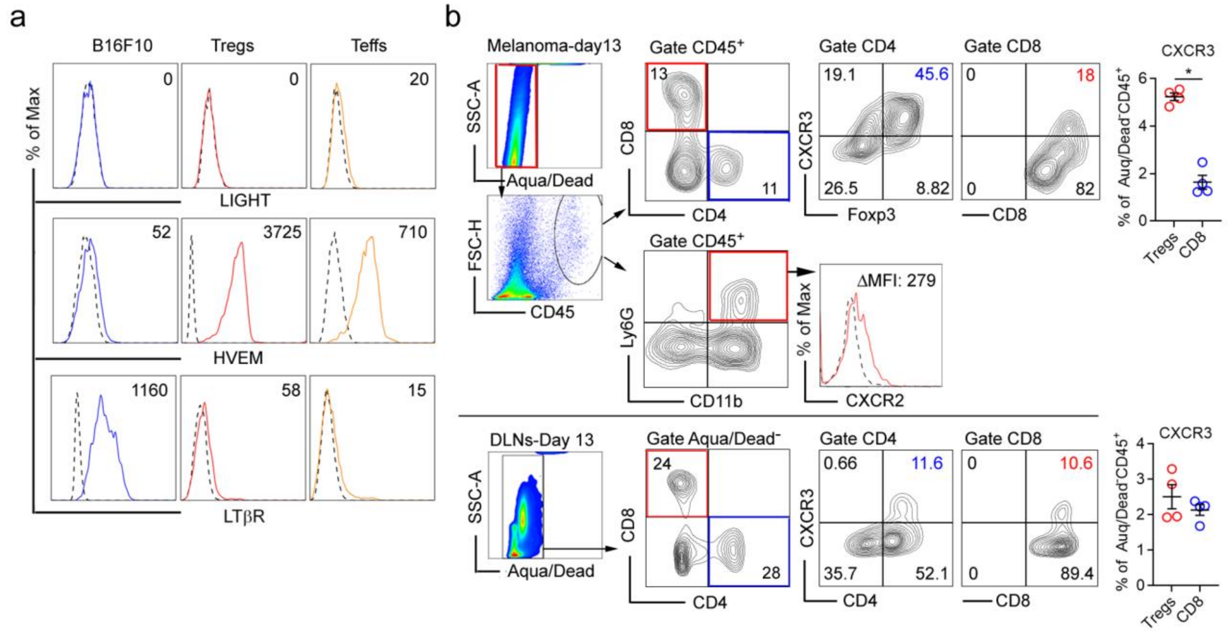


(a) Flow cytometry analysis of LIGHT, HVEM, and LTβR on B16F10, Tregs, and Teffs. ΔMFI shown. (b) Expression of CXCR2 on MDSCs; CXCR3 on Tregs and CD8 in melanoma TILs and dLNs in day 13. ΔMFI shown. Data represent 2 independent experiments (WT B16F10/WT C57BL/6 mice from Fig.4d and Fig.6j). Mean ± SEM. *$p < 0.05$ by one-way ANOVA.


1 Wu, S. Z. *et al.* Cryopreservation of human cancers conserves tumour heterogeneity for single-cell multi-omics analysis. *Genome Med* **13**, 81, doi:10.1186/s13073-021-00885-z (2021).

2 Zilionis, R. *et al.* Single-Cell Transcriptomics of Human and Mouse Lung Cancers Reveals Conserved Myeloid Populations across Individuals and Species. *Immunity* **50**, 1317-1334 e1310, doi:10.1016/j.immuni.2019.03.009 (2019).

3 Wu, S. Z. *et al.* A single-cell and spatially resolved atlas of human breast cancers. *Nat Genet* **53**, 1334-1347, doi:10.1038/s41588-021-00911-1 (2021).

4 He, M. X. *et al.* Transcriptional mediators of treatment resistance in lethal prostate cancer. *Nat Med* **27**, 426-433, doi:10.1038/s41591-021-01244-6 (2021).
